# Mechanochemical tuning of a kinesin motor essential for malaria parasite transmission

**DOI:** 10.1101/2022.02.11.480087

**Authors:** Tianyang Liu, Fiona Shilliday, Alexander D. Cook, Rita Tewari, Colin J. Sutherland, Anthony. J. Roberts, Carolyn A. Moores

**Affiliations:** Institute of Structural and Molecular Biology, Birkbeck College, London WC1E 7HX, UK; Department of Biochemistry, University of Oxford, Oxford OX1 3QU; School of Life Sciences, University of Nottingham, Nottingham, UK; Department of Infection Biology, Faculty of Infectious & Tropical Diseases, London School of Hygiene & Tropical Medicine

**Keywords:** ATPase, cryo-EM, flagella, kinesin, malaria, microtubule, motor

## Abstract

Kinesins are a superfamily of molecular motors that undertake ATP-dependent microtubule-based movement or regulate microtubule dynamics. *Plasmodium* species, which cause malaria and kill hundreds of thousands annually, encode 8-9 kinesins in their genomes. Of these, two – kinesin-8B and kinesin-8X -are canonically classified as kinesin-8s, which in other eukaryotes typically modulate microtubule dynamics. Unexpectedly, *Plasmodium* kinesin-8B is required for development of the flagellated male gamete in the mosquito host, and its absence completely blocks parasite transmission. To understand the molecular basis of kinesin-8B’s essential role, we characterised the *in vitro* properties of the kinesin-8B motor domains from *P. berghei* and *P. falciparum*. Both motors drive plus-end directed ATP-dependent microtubule gliding, but also catalyse ATP-dependent microtubule depolymerisation. We determined the microtubule-bound structures of these motors using cryo-electron microscopy. *P. berghei* and *P. falciparum* kinesin-8B exhibit a very similar mode of microtubule interaction, in which *Plasmodium*-distinct sequences at the microtubule-kinesin interface influence motor function. Intriguingly, however, *P. berghei* kinesin-8B exhibits a non-canonical structural response to ATP analogue binding such that neck linker docking is not induced. Nevertheless, the neck linker region is absolutely required for motility and depolymerisation activities of these motors. Taken together, these data suggest that the mechanochemistry of *Plasmodium* kinesin-8Bs has been functionally tuned to efficiently contribute to flagella formation.

## INTRODUCTION

Malaria – of which there were 241 million cases globally and 627,000 deaths in 2020 (https://www.who.int/publications/i/item/9789240040496) – is caused by apicomplexan *Plasmodium* parasites. *Plasmodium spp*. are obligate intracellular parasites with a complex life cycle that alternates between mammalian hosts and mosquito vectors. The microtubule (MT) cytoskeleton plays a number of important roles throughout this life cycle, including formation of the mitotic/meiotic spindles during the several replicative stages^1^, during invasion of and egress from host cells and tissues^2^, and in forming the motile flagella in male gametes^3^. Given this diversity of functions, precise regulation of MT dynamics and organisation by cellular factors is absolutely essential for parasite survival. In particular, the flagella-driven motility of male gametes, which develop from male gametocytes in the mosquito gut immediately on ingestion of a blood meal, is required to fertilise female gametes for onward progression of the life cycle. If male gamete motility is compromised, parasite transmission is blocked^3^. Therefore, understanding the molecular processes involved in male gamete development and flagella formation is of fundamental interest, and may offer new avenues for development of disease control^4^.

Kinesin-8B - a member of the kinesin superfamily of ATP-driven, MT-based molecular motors^5^- is required for flagella formation in *P. berghei* male gametes, and its knockout completely disrupts parasite transmission^6,7^. Kinesin-8s are one of the fifteen kinesin families identified across apicomplexa^8^ that, together with kinesin-13s, are important regulators of MT dynamics^9^. Kinesin-8s are phylogenetically subclassified into kinesin-8As (e.g. mammalian KIF18A, KIF18B, *S. cerevisiae* Kip3), kinesin-8Bs (mammalian KIF19A) and kinesin-8Xs^8^ (*Plasmodium* kinesin-8X^10^). There are up to 9 kinesin genes in *Plasmodium spp*.^10^ and, intriguingly, with genes encoding one kinesin-13 and two kinesin-8 isoforms (kinesin-8B and kinesin-8X), at least one third of *Plasmodium* kinesins are potential regulators of MT dynamics^8^. This likely reflects the requirement for frequent and often rapid remodelling of the MT cytoskeleton during the parasite life cycle.

Although eukaryote-wide kinesin families have been distinguished based on phylogenetics^8,10^, we don’t yet know if individual families such as kinesin-8s have conserved molecular activities and function. Previously characterised kinesin-8s (primarily from mammals and yeast) move towards the plus-ends of MTs and regulate dynamics of these ends on arrival^9^. While lattice-based kinesin-8 movement requires the ATPase activity of the motor domains, ATP binding but not necessarily hydrolysis appears to be required for MT end regulation^11^. Kinesin-8s also exhibit MT depolymerisation activity, but their depolymerisation mechanism and the extent of conservation of this activity is not currently clear^12-15^. The best understood function of kinesin-8s is regulation of spindle MT dynamics during chromosome alignment^16^. *P. berghei* kinesin-8X is spindle-associated in the mosquito stages of the parasite life cycle and is needed for oocyst development and sporozoite formation. This motor exhibits plus-end directed motility and MT depolymerisation activity, supporting a classical role for this motor in regulating spindle dynamics^10^.

What are the molecular properties of *Plasmodium* kinesin-8B that support its essential function in flagella formation? To answer this question, we studied the activities and structures of kinesin-8B motor domains from both *P. berghei* and *P. falciparum*, referred to here as *Pb*kinesin-8B-MD and *Pf*kinesin-8B-MD (Fig. 1a). These monomeric constructs share 88% sequence identity (Supplementary Fig. 1a), and we show that they both drive MT plus-end directed motility. *Pb*kinesin-8B-MD and *Pf*kinesin-8B-MD also catalyse ATPase-dependent MT depolymerisation, demonstrating their multi-tasking capabilities with respect to MTs. The structures of the motors bound to MTs determined using cryo-electron microscopy (cryo-EM) exhibit a canonical kinesin fold. Our reconstructions visualise distinct regions of MT interaction, mutation of which perturbs motor activity. However, *Pb*kinesin-8B-MD exhibits a non-canonical structural response to ATP analogue binding, with no nucleotide binding site closure or neck linker docking observed. We nevertheless also find that removal of the neck-linker region abolishes both MT motility and depolymerisation activities. By comparing the structural and biochemical properties of *Pb*kinesin-8B-MD to those of previously characterised kinesin regulators of MT dynamics, we conclude that *Plasmodium* kinesin-8Bs exhibit a blend of canonical properties characteristic of both kinesin-8s and non-motile kinesin-13s. Together, our data demonstrate how kinesin mechanochemistry has been tuned for particular cellular roles across kinesin subfamilies and in evolutionary divergent organisms.

**Figure 1.**
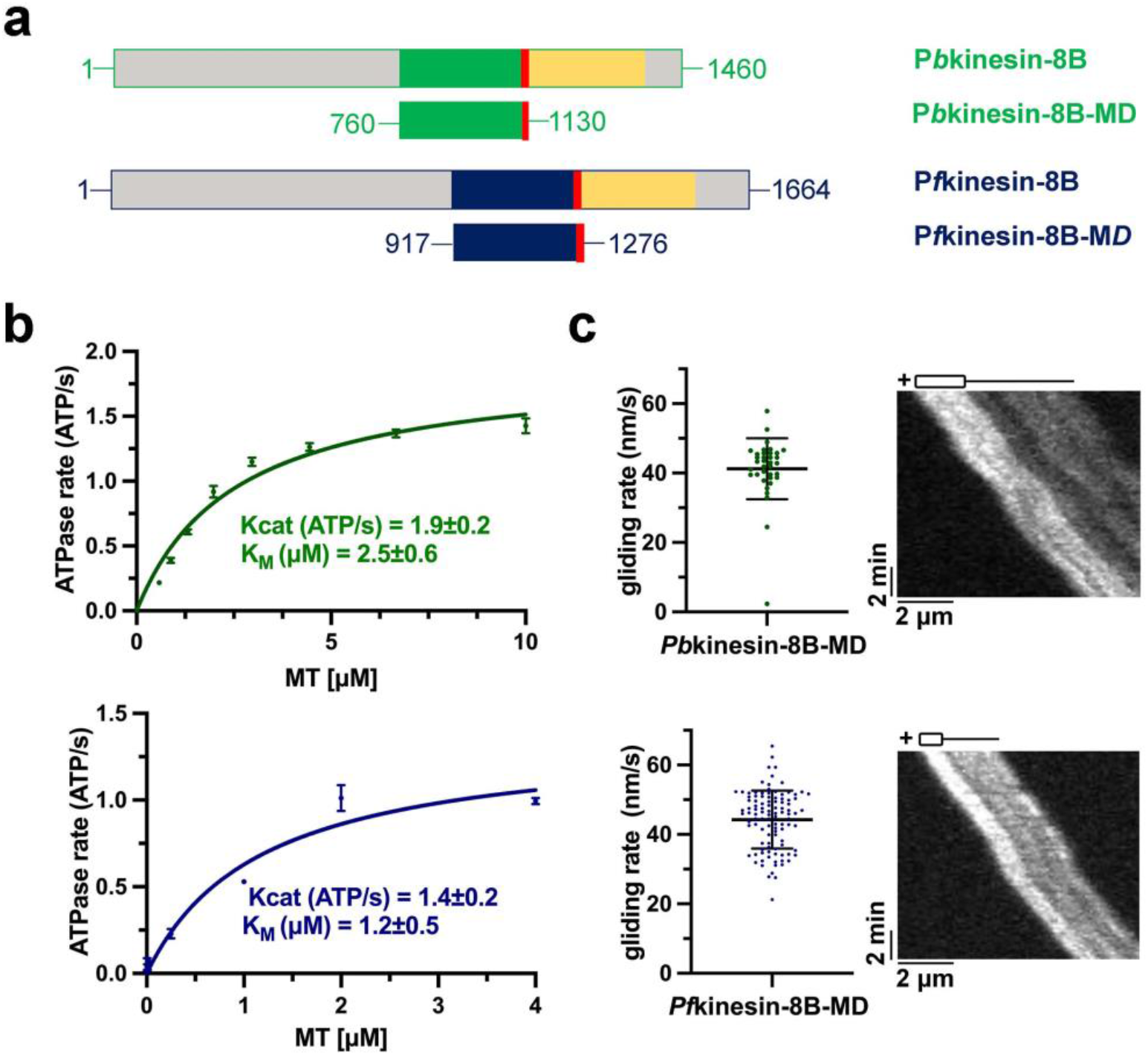
*Plasmodium* kinesin-8B motor domains are MT-dependent ATPases and drive plus-end directed MT gliding. a) Schematic representation of full-length *Pb*kinesin-8B (PBANKA_020270, top) and *Pf*kinesin-8B (PF3D7_0111000, bottom) domain organisation, indicating the relationship with the MD construct (the motor domain plus NL sequence); motor domains are coloured green (*Pb*kinesin-8B) and blue (*Pf*kinesin-8B), neck linkers are red and coiled-coil regions are yellow. b) Both *Pb*kinesin-8B-MD (top - green) and *Pf*kinesin-8B-MD (bottom – blue) exhibit MT stimulated ATPase activity (*Pb*:GMPCPP-MT, *Pf*:paclitaxel-stabilized MT). ATPase assay data (n=3 for each point, mean ± SD) were fitted using the Michaelis-Menten equation, from which the Kcat and K_M_ were calculated in Prism 9. c) Both *Pb*kinesin-8B-MD (top - green) and *Pf*kinesin-8B-MD (bottom – blue) exhibit MT-plus end directed gliding activity. For *Pb*kinesin-8B-MD, the velocity = 41.3± 8.8 nm/s (mean ± SD; n = 36), and for *Pf*kinesin-8B-MD = 44.3 ± 8.4 nm/s (mean ± SD; n = 104). Data are plotted on the left, while the representative TIRF-M kymographs on the right shows gliding of a single polarized MT consistent with plus-end directed motility; MT schematic above.

## RESULTS

### *Pb*kinesin-8B-MD and *Pf*kinesin-8B-MD have MT-stimulated ATPase and motility activities

To investigate the molecular properties of *Plasmodium* kinesin-8Bs, we expressed and purified *Pb*kinesin-8B-MD and *Pf*kinesin-8B-MD (Fig. 1a; Supplementary Fig. 1b), and measured their steady state MT-stimulated ATPase activities (Fig. 1b). The activities of each of these constructs were comparable: Kcat = 1.9±0.2 ATP/s and Km = 2.5±0.6 μM for *Pb*kinesin-8B-MD, while Kcat = 1.4±0.2ATP/s and Km = 1.2±0.5μM for *Pf*kinesin-8B-MD.

To begin to understand how the ATPase activity of these motors is harnessed, we investigated their behaviour in a multi-motor gliding assay using TIRF microscopy (TIRF-M), in which motors are attached to the assay coverslip and labelled, stabilised MTs are flowed into the assay cell. Both kinesins generated MT movement, with average velocities of *Pb*kinesin-8B-MD = 41.3 ± 8.8 nm/s and *Pf*kinesin-8B-MD = 44.3 ± 8.4 nm/s (Fig.1c, left). Using polarised GMPCPP MTs, we observed that this gliding activity was plus-end directed for both motors (Fig.1c, right). These data establish that the biochemical properties of these parasite kinesin-8B constructs are conserved, and that they are capable of driving ATP-dependent plus-end directed motility along MTs.

### *Pb*kinesin-8B-MD and *Pf*kinesin-8B-MD are MT depolymerases

We also used TIRF-M to investigate the influence of *Pb*kinesin-8B-MD and *Pf*kinesin-8B-MD on MT ends. In this assay, we incubated unlabelled motor protein with tethered, labelled, paclitaxel-stabilised MTs and monitored MT length. Both kinesin-8B constructs cause MT shortening in the presence of ATP or the non-hydrolysable ATP analogue AMPPNP (Fig. 2a). In all cases, shortening is observed at both MT ends showing that the MT depolymerisation activity of *Pb*kinesin-8B-MD and *Pf*kinesin-8B-MD is not restricted by the plus-end directed motility of these constructs (Fig. 1c). Our data are consistent with depolymerisation occurring as a result of monomeric motors encountering both MT ends by diffusion from solution. It also supports previous observations that dimeric motor-mediated stepping along the MT lattice is not required for kinesin-8-mediated depolymerisation at MT ends^11,17,18^.

**Figure 2.**
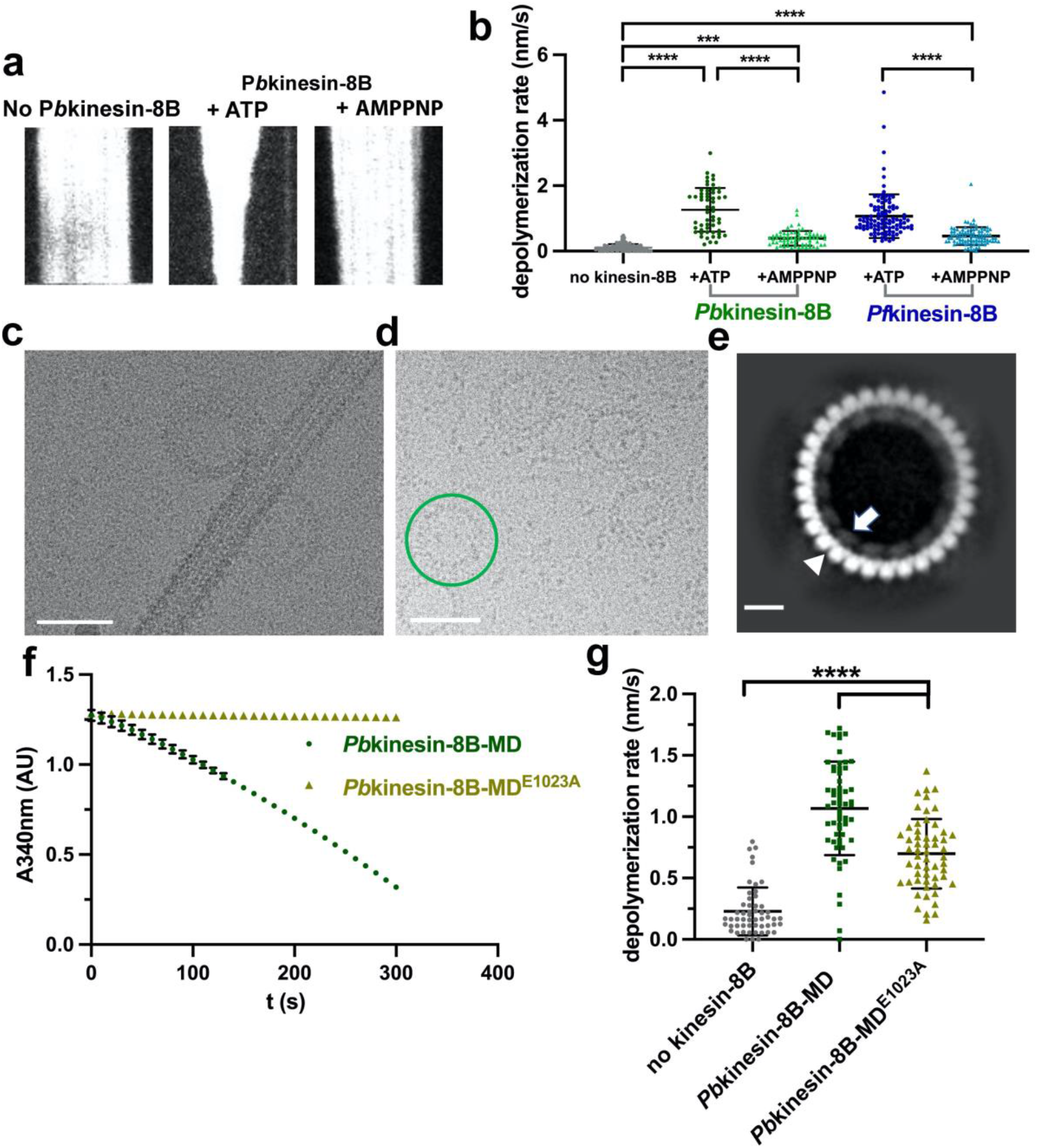
*Plasmodium* kinesin-8B motor domains are MT depolymerases. a) Representative TIRF-M kymographs of *Pb*kinesin-8B-MD depolymerising paclitaxel-stabilised MTs in the presence of ATP (middle) and AMPPNP (right). Depolymerisation occurs at both MT ends in both conditions; b) MT depolymerisation rate (nm/s) for *Pb*kinesin-8B-MD and *Pf*kinesin-8B-MD in the presence of ATP or AMPPNP compared to the no kinesin control. Error bars represent the mean ±SD and individual measurements are also plotted. Ordinary one-way ANOVA was performed in Prism. Significance values are displayed as asterisks, ****, p-values <0.0001; ***, p-value < 0.001. n_no-kinesin-8B_= 112 ends, n_PbKin-8B-MD ,ATP_ = 56 ends, n_PbKin-8B-MD ,AMP_ = 68 ends, n_PfKin8B-MD ,ATP_ = 103 ends, n_PfKin-8B-MD ,AMP_ = 92 ends. c) Cryo-EM image showing protofilaments peeling from MT wall and forming ring-like structures. d) Cryo-EM image showing ring structures formed by incubating tubulin and *Pb*kinesin-8B-MD in the presence of AMPPNP. A representative ring structure is highlighted with green circle. Scale bar in c, d = 50 nm. e) Representative 2D class average of AMPPNP-dependent *Pb*kinesin-8B-MD-induced tubulin ring structure; the outer ring is formed by curved αβ-tubulin dimers (white arrowhead), while and inner ring is formed by individual *Pb*kinesin-8B-MD density (white arrow). Scale bar = 10nm. *f) Pb*kinesin-8B-MD^E1023A^ (olive green) does not exhibit MT-stimulated ATPase activity in the enzyme-coupled assay used for measuring ATPase activity (n=3 for each point), while WT *Pb*kinesin-8B-MD (dark green) induces a decrease in NADH absorbance (340 nm), both in the presence of 1μM MTs. g) MT depolymerisation rate (nm/s) for *Pb*kinesin-8B-MD and *Pb*kinesin-8B-MD^E1023A^ in the presence of ATP compared to the no kinesin control. Error bars represent the mean ± SD and individual measurements are also plotted. Ordinary one-way ANOVA was performed in Prism. Significance values are displayed as asterisks, ****, p-values <0.0001; n_no kinesin-8B_= 54 ends, n_PbKin-8B-MD ,ATP_ = 60 ends, n_PbKin-8B-MD_^E1023A^ ,_ATP_ = 56 ends.

In the presence of ATP, the average depolymerisation rate by *Pb*kinesin-8B-MD is 1.3 ± 0.7 nm/s and by *Pf*kinesin-8B-MD is 1.1 ± 0.7 nm/s (Fig. 2b). In the presence of the non-hydrolysable ATP analogue, AMPPNP, the depolymerisation is slower -by *Pb*kinesin-8B-MD it is 0.4 ± 0.2 nm/s and 0.5 ± 0.3 nm/s by *Pf*kinesin-8B-MD (Fig. 2b). Faster depolymerisation in the presence of ATP compared to AMPPNP demonstrates that ATPase cycle turnover of *Pb*kinesin-8B-MD and *Pf*kinesin-8B-MD is coupled to catalytic MT depolymerisation. During ATP turnover, these motors can interact with MT ends, induce tubulin release, themselves release from this tubulin and thus be recycled for further depolymerisation. The observation that some depolymerisation occurs in the presence of AMPPNP shows that the ATP-binding step of the motor’s ATPase cycle can be sufficient for tubulin release. However, the slower overall depolymerisation without ATP hydrolysis at the same motor concentration suggests the motors could be trapped on depolymerisation products. Consistent with this, we observed formation of tubulin rings and peeling protofilaments when *Pb*kinesin-8B-MD was incubated with stabilised MTs and AMPPNP but not ATP (Fig. 2c). Such curved structures do not form from stabilised MTs in the absence of *Pb*kinesin-8B-MD. Similar but more plentiful rings and spirals were also observed on incubation of *Pb*kinesin-8B-MD with AMPPNP and unpolymerized tubulin (Fig. 2d). Although these oligomers are flexible and heterogeneous, 2D image analysis showed that *Pb*kinesin-8B-MD molecules bind to curved tubulin dimers around the inner circumference of these rings (Fig. 2e). These observations demonstrate that the ATP-binding step of malaria kinesin-8B motors can induce or stabilise a bent tubulin conformation which drives tubulin release from MT ends, an activity that is not typical of other kinesin-8s^11,18,19^.

To further investigate the relationship between MT depolymerisation activity and nucleotide hydrolysis, we prepared a *Pb*kinesin-8B-MD ATPase inactive mutant, in which the Glu residue in the conserved switch 2 motif (DXXGXE) is mutated to Ala (*Pb*kinesin-8B-MD ^E1023A^)^20,21^. As expected, *Pb*kinesin-8B-MD ^E1023A^ exhibited no ATPase activity (Fig. 2f). This mutant was nevertheless able to depolymerise stabilised MTs in the presence of ATP, with a mean rate of 0.7 ± 0.3 nm/s (Fig.2g). This is 65% of the WT +ATP rate (one-way ANOVA; p<0.0001), compared to 31% of the WT +ATP rate observed in the presence of AMPPNP. (Fig. 2b; one-way ANOVA; p < 0.001). This further supports the idea that while ATP turnover by *Plasmodium* kinesin-8Bs is not essential for MT depolymerisation, it supports catalytic MT depolymerisation by these motors.

Individual tubulin dimers have been suggested to share some structural properties with tubulins located at MT ends, so we also measured the tubulin-stimulated ATPase activity of *Pb*kinesin-8B-MD (Supplementary Fig. 2). Although tubulin was found to stimulate the motor ATPase to some extent, motor turnover is much slower than with MTs (Kcat = 0.1 ±0.0 ATP/s) and its interaction is also weaker (Km = 7.6 ±2.4 μM). The ratio of MT-stimulated ATPase Kcat compared to tubulin is thus much higher (19x) for *Pb*kinesin-8B-MD than mammalian kinesin-8s (KIF18A_MD ratio = 5.0; KIF19A_MD ratio = 1.8;^17,19^). Although tubulin may not optimally mimic the configuration of the MT end substrate for *Plasmodium* kinesin-8 depolymerisation activity, these data suggest that the lattice-based ATPase activity of the parasite motor domains dominates compared to the depolymerisation activity at MT ends.

### Nucleotide-dependent structures of lattice-bound *Plasmodium* kinesin-8Bs

To investigate the mechanistic basis for these activities, we used cryo-EM to determine the structures of *Plasmodium* kinesin-8Bs bound to MTs (Fig. 3). MT-bound complexes of *Pb*kinesin-8B-MD in 2 different nucleotide states – no nucleotide (NN) and AMPPNP -were imaged and their structures determined to overall resolutions of 4.3 and 3.3 Å respectively (Table 1; Supplementary Fig. 3). Resolutions in the kinesin motor domain of each reconstruction ranged between 4 and 8 Å (Supplementary Fig. 3). MT-bound complexes of *Pf*kinesin-8B-MD in the absence of nucleotide (NN) were also imaged and used to calculate a reconstruction with an overall resolution of 4.1 Å with resolution in the kinesin motor domain of 4-5 Å (Supplementary Fig. 4a). To facilitate interpretation of these structures, we built molecular models of the motor-MT complexes (Table 2).

**Table 1.** Cryo-EM data collection, 3D image processing statistics.

| Data collection and reconstruction | Pbkinesin-8B-MD-NN | Pbkinesin-8B-MD-AMPPNP | Pfkinesin-8B-MD-NN |
| --- | --- | --- | --- |
| Grid type | C-Flat 2/2-4C | C-Flat 2/2-4C | C-Flat 2/2-4C |
| Microscope | Polara | Krios | Krios |
| Detector and mode | K2 counting mode | K2 counting mode | K2 counting mode |
| Collection software | Serial EM | EPU | EPU |
| Magnification | 160K | 130K | 130K |
| Voltage(Kv) | 300 | 300 | 300 |
| Electron exposure (e<sup>-</sup>/ Å<sup>2</sup>) | 50.81 | 47.11 | 46.50 |
| Exposure time(s) | 15 | 8 | 8 |
| Dose rate (e<sup>-</sup>/pixel/s) | 6.54 | 6.80 | 6.71 |
| Total frame | 50 | 32 | 32 |
| Fraction dose (e<sup>-</sup>/ Å<sup>2</sup>) | 1.02 | 1.47 | 1.45 |
| Defocus range(um) | -0.5 to -2.5 | -0.5 to -2.5 | -0.5 to -2.5 |
| Pixel size(Å) | 1.35 | 1.05 | 1.05 |
| Particle number for final reconstruction | 196084 | 87907 | 205687 |
| Map resolution (Å,FSC0.143) | 4.3 | 3.3 | 4.1 |
| Local resolution range(Å) | 4.2-7.6 | 3.3-6.5 | 3.6-5 |
| B factor | -120 | -20 | -20 |

**Table 2.**
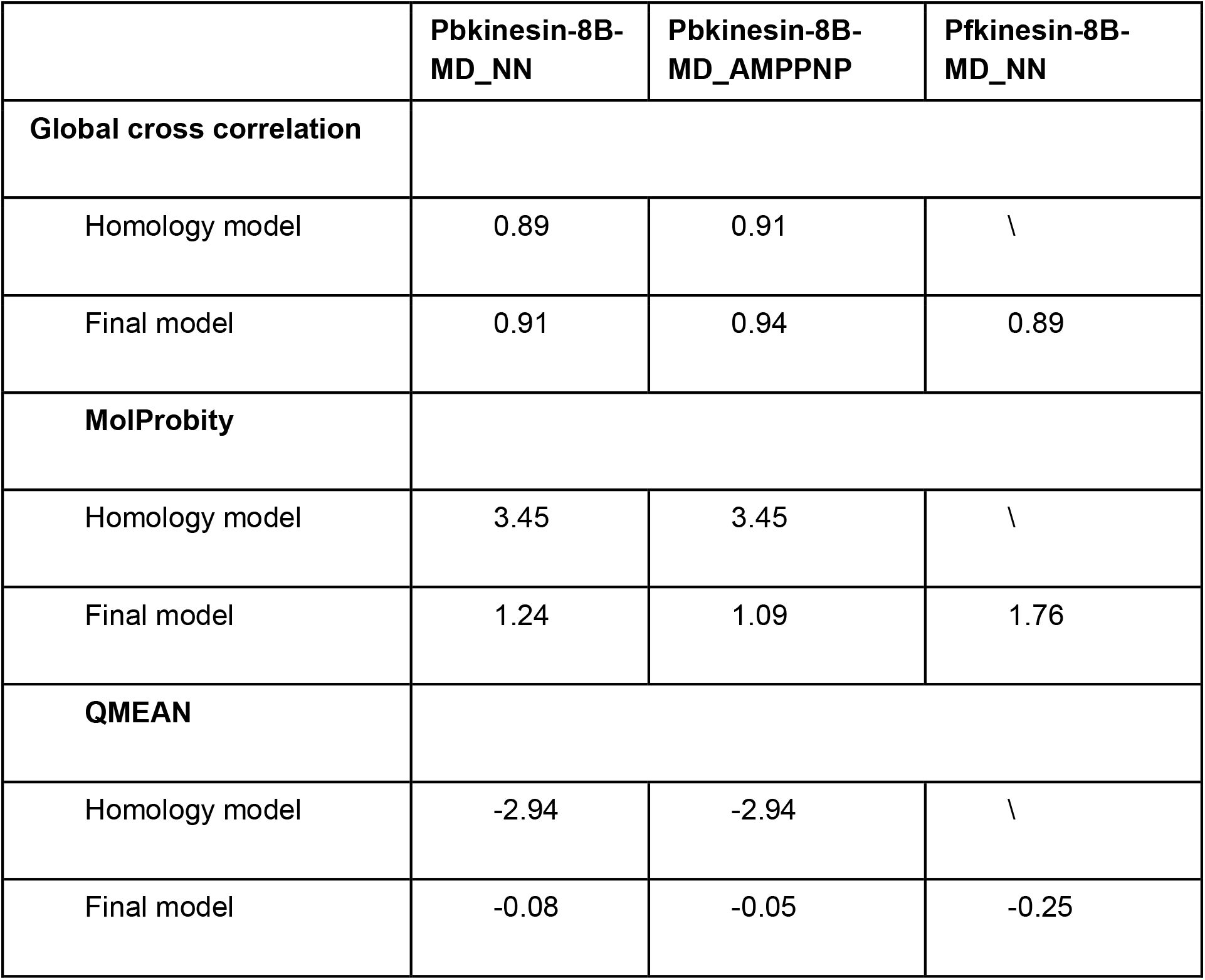
Model building statistics.

|  | **Pbkinesin-8B-MD_NN** | **Pbkinesin-8B-MD_AMPPNP** | **Pfkinesin-8B-MD_NN** |
| --- | --- | --- | --- |
| **Global cross correlation** |  |  |  |
| Homology model | 0.89 | 0.91 | \ |
| Final model | 0.91 | 0.94 | 0.89 |
| **MolProbity** |  |  |  |
| Homology model | 3.45 | 3.45 | \ |
| Final model | 1.24 | 1.09 | 1.76 |
| **QMEAN** |  |  |  |
| Homology model | -2.94 | -2.94 | \ |
| Final model | -0.08 | -0.05 | -0.25 |

**Figure 3.**
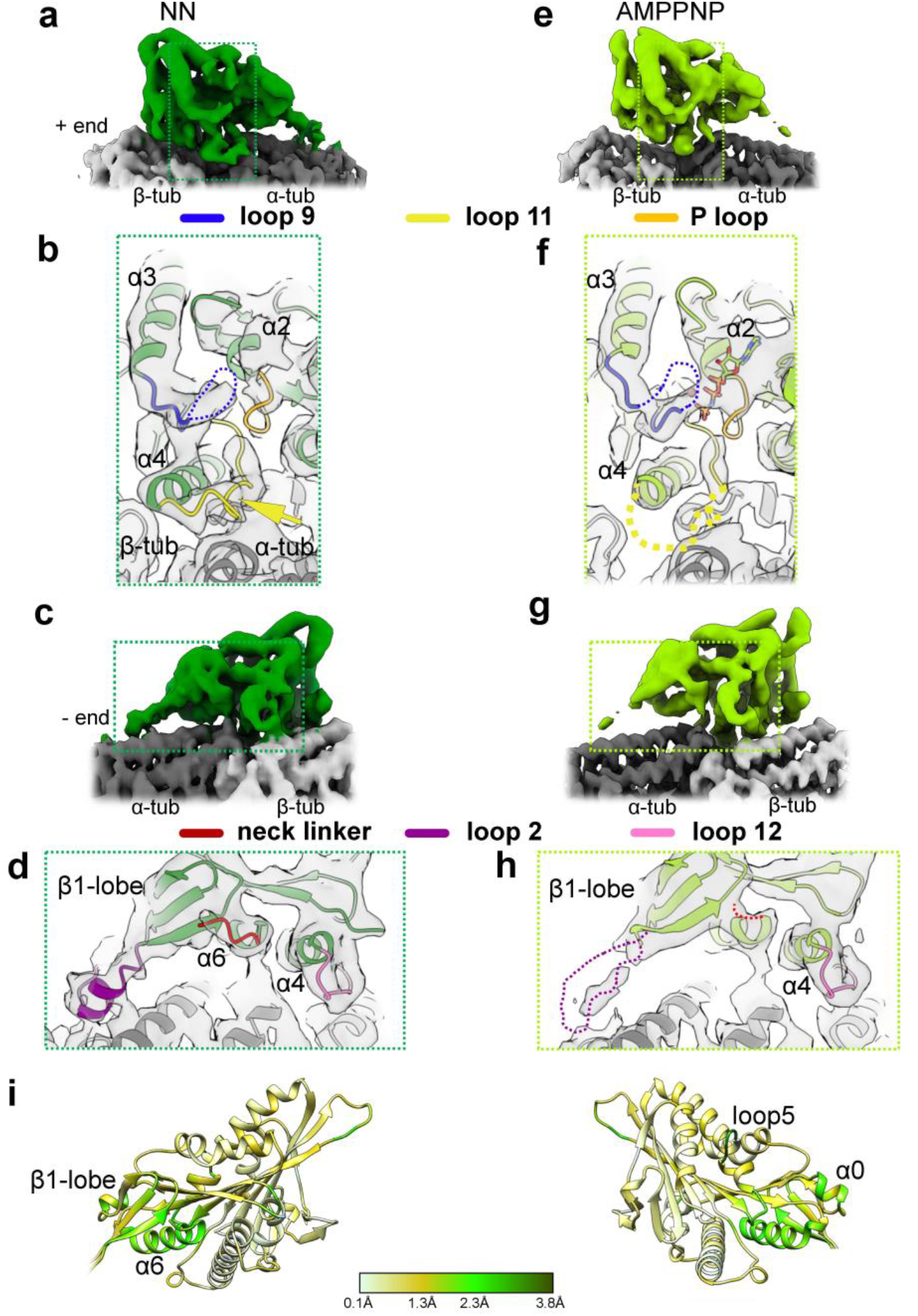
Cryo-EM reconstructions of MT-bound *Pb*kinesin-8B-MD. a) Asymmetric unit of MT-bound NN *Pb*kinesin-8B-MD depicted as solid surface representation and viewed towards the NBS. *Pb*kinesin-8B-MD-NN density is coloured in dark green, α- and β-tubulin are coloured in dark and light grey, respectively; region around NBS depicted in (b) is boxed. b) Zoom-in view of the NBS of NN *Pb*kinesin-8B-MD with docked model, showing the contact formed between the helical turn (arrow) in loop11 (yellow) and α-tubulin, the P-loop (orange) in the empty NBS, and the density corresponding to flexible-appearing loop 9 (blue). *Pb*kinesin-8B-MD-NN model is coloured in dark green and α- and β-tubulin are coloured in dark and light grey, respectively. c) MT-bound NN *Pb*kinesin-8B-MD depicted as solid surface representation viewed towards the neck linker region. *Pb*kinesin-8B-MD-NN density is coloured in dark green, α- and β-tubulin are coloured in dark and light grey, respectively; region around the neck linker depicted in (d) is boxed. d) Zoom-in view of the neck linker region of NN *Pb*kinesin-8B-MD with docked model, showing density corresponding to loop12 (fuchsia) at the C-terminus of helix-α4 that contacts β-tubulin, adjacent to which is density corresponding to the N-terminal end of the neck linker (red), which is directed towards the MT minus end. e) Asymmetric unit of MT-bound AMPPNP *Pb*kinesin-8B-MD depicted as solid surface representation and viewed towards the NBS. *Pb*kinesin-8B-MD-AMPPNP density is coloured in light green, α- and β-tubulin are coloured in dark and light grey, respectively; region around NBS depicted in (f) is boxed. f) Zoom-in view of the NBS of AMPPNP *Pb*kinesin-8B-MD with docked model, showing weaker loop 11 density (dashed yellow line), the P-loop (orange) adjacent to density corresponding to AMPPNP in the NBS and density corresponding to flexible-appearing loop 9 (dashed blue line). AMPPNP *Pb*kinesin-8B-MD model is coloured in light green and α- and β-tubulin are coloured in dark and light grey, respectively. g) MT-bound AMPPNP *Pb*kinesin-8B-MD depicted as solid surface representation viewed towards the neck linker region. *Pb*kinesin-8B-MD-AMPPNP density is coloured in light green, α- and β-tubulin are coloured in dark and light grey, respectively; region around the neck linker depicted in (h) is boxed. h) Zoom-in view of the neck linker region of AMPPNP *Pb*kinesin-8B-MD with docked model, showing density corresponding to loop12 (fuchsia) at the C-terminus of helix-α4 that contacts β-tubulin, and weaker neck linker density (red dotted line), directed towards the MT minus end. i) Cα RMSD in Å of *Pb*kinesin-8B-MD NN compared to AMPPNP models aligned on helix-α4 of *Pb*kinesin-8B-MD, depicted on the NN model; the small range of RMSD observed illustrated that only minor structural changes are detected when AMPPNP binds.

All the reconstructions show that *Pb*kinesin-8B-MD and *Pf*kinesin-8B-MD contact a single tubulin dimer in the MT lattice, with motor binding centred on the intradimer tubulin dimer interface (Fig. 3;Supplementary Fig. 4b). In the NN structure of *Pb*kinesin-8B-MD, density corresponding to nucleotide is indeed absent from the nucleotide binding site (NBS) (Fig. 3a, b). The conserved nucleotide binding loops – P-loop, loop 9 (containing the switch I motif) and loop 11 (containing the switch II motif) – adopt a canonical conformation previously described for the NN state of a number of other plus-end directed kinesins, including human KIF18A^18^. In this conformation, density corresponding to the P-loop is visible in the empty NBS, while loop11 is retracted from the NBS. The C-terminal end of loop 11 adopts a turn and interacts with α-tubulin before it leads into helix-α4, a major contact point with the MT surface. Density corresponding to loop 9 is visible between the P-loop and loop 11 but is poorly defined. Adjustment of the reconstruction density threshold reveals some evidence of connectivity between loop9 and both the P-loop and the helical turn of loop 11, supporting the idea that loop 9 is partially flexible in the absence of bound nucleotide. Consistent with this conformation of the NBS, on the opposite side of the kinesin motor domain, helix-α6 abuts the C-terminal end of helix-α4 and density corresponding to the beginning of the C-terminal neck linker peptide of *Pb*kinesin-8B-MD is visible protruding towards the MT minus end and adjacent to the β1-lobe (Fig. 3c, d). The N-terminal peptide of the motor can also be visualised protruding in the opposite direction towards the MT plus end. The NN *Pf*kinesin-8B-MD reconstruction is very similar to that of *Pb*kinesin-8B-MD (Supplementary Fig. 4b,c), including the empty NBS and undocked neck linker. Overall, the configuration of MT-bound NN *Plasmodium* kinesin-8Bs in these conserved parts of their motor domains are similar to a number of NN states of other plus-end directed motors^18,22-24^.

Surprisingly, the structure of MT-bound *Pb*kinesin-8B-MD in the presence of AMPPNP is overall similar to that of the NN state (Fig. 3e-h). While density corresponding to bound nucleotide is clearly present (Supplementary Fig. 5), the overall open configuration of the NBS is very similar to that in the motor’s NN conformation (Fig. 3e, f). Consistent with this, helix-α6 again abuts the C-terminal end of helix-α4 and no neck linker docking is observed (Fig. 3g,h). However, differences in EM density are visible between the 2 nucleotide states at lower thresholds, which show that density corresponding to a number of loops around the motor domain - including loop 2, loop 9, loop 11 and the neck linker - are more flexible in the presence of bound AMPPNP and, as a consequence, were not included in the AMPPNP model (Fig. 3f, h dashed lines). Because this variation in density is not attributable to resolution differences, we conclude that it reflects small structural adjustments of the motor domain to the bound nucleotide, but these are not converted into larger conformational changes. As a result, overlay of the NN and AMPPNP models of *Pb*kinesin-8B-MD (aligned on helix-α4) shows only minor structural variations around the NBS, β1-lobe and in helix-α6 (Fig. 3i). Such changes are very small compared to the changes that have been observed in the NN-AMPPNP transition in other plus ended kinesins^23-26^.

### MT-binding interface of *Plasmodium* kinesin-8Bs and distinct contributions of interface regions to *Pb*kinesin-8B-MD function

The interaction between *Plasmodium* kinesin-8Bs and α- and β-tubulin is centred on helix-α4 (Fig. 3, Fig. 4a, Supplementary Fig. 4). Contacts are also formed with β-tubulin by the C-terminal part of loop 12, helix-α5 and β2-lobe/loop 8, and between α-tubulin and helix-α6 (Fig. 4a). The MT interaction in all these regions is not detectably different between the *Pb*kinesin-8B-MD NN and AMPPNP reconstructions (Supplementary Fig. 6a). These elements are well conserved points of MT contact in kinesins from different families ^22,24-27^,, although loop 12 often exhibits family-specific insertions, including in *Plasmodium* kinesin-8Bs (Fig. 4b)^23,28^. In both *Plasmodium* kinesin-8Bs NN reconstructions, there is an additional connection between α-tubulin and loop 2, which protrudes from the β1-lobe of the motor domain and appears to adopt a partially helical configuration (Fig. 3a, d, Fig. 4a). In the *Pb*kinesin-8B-MD AMPPNP reconstruction, the density corresponding to loop 2 is less distinct due to the above described motor domain flexibility, although at more inclusive density thresholds, connectivity with the MT surface is also visible (Supplementary Fig. 6b). *Plasmodium* kinesin-8B loop 2 is the same length as, and relatively well-conserved compared to, loop 2 in mammalian kinesin-8B KIF19A (Supplementary Fig. 7a,b)^19^, although shorter compared to loop 2 in the mammalian kinesin-8A KIF18A (Supplementary Fig. 7a). Most kinesin-8 proteins so far characterised form an additional MT contact via loop 2^18,19^, a characteristic we now show *Plasmodium* kinesin-8Bs also share.

**Figure 4.**
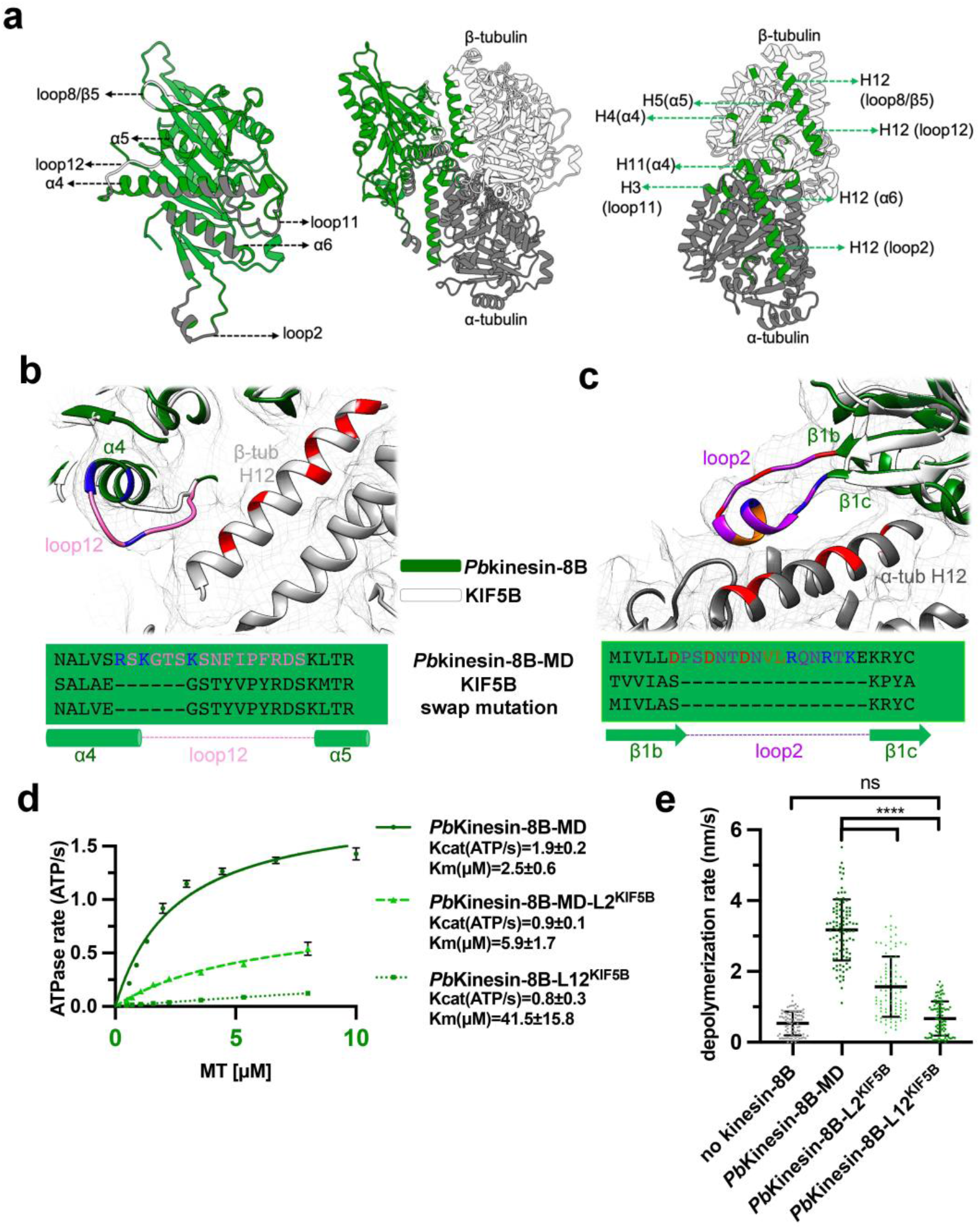
The MT binding interface of *Pb*kinesin-8B-MD and contributions to motor function. a) Middle, ribbon depiction of the *Pb*kinesin-8B-MD and tubulin dimer NN state model, with *Pb*kinesin-8B-MD in green, α-tubulin in dark grey and β-tubulin in light grey; left, zoomed view of the *Pb*kinesin-8B-MD MT binding surface coloured according to contacts with α-tubulin (dark grey) and β-tubulin (light grey); right, MT footprint of *Pb*kinesin-8B-MD on α- and β-tubulin indicated in dark green (tubulin residues <5Å distance from the bound motor). Labels indicate the specific contacting secondary structure elements in tubulin dimer and *Pb*kinesin- 8B-MD(in bracket). b) Structural alignment of the *Pb*Kinesin-8B-MD model (green) and KIF5B motor domain model (PDB 6ojq, white), focusing on the loop12, with the *Pb*Kinesin-8B-MD NN cryo-EM density shown in mesh representation. A sequence alignment of this region, and the swap mutant, is depicted below. *Pb*Kinesin-8B-MD loop12 is coloured purple with positively charged residues coloured blue. Negatively charged residues in the adjacent H12 of β-tubulin are coloured red; c) Structural alignment of the *Pb*Kinesin-8B-MD model (green) and KIF5B motor domain model (PDB 6ojq, white), focusing on loop2, with the *Pb*Kinesin-8B-MD NN cryo-EM density shown in mesh. A sequence alignment of this region, and the sequence of the swap mutant, is depicted below. *Pb*Kinesin-8B-MD loop2 is coloured pink with positively charged residues coloured blue, negatively charged residues coloured red and hydrophobic residues coloured orange. Negatively charged residues in the adjacent H12 of α-tubulin are also coloured red, indicating the potential electrostatic interactions between loop2 and the MT surface. d) GMPCPP-MT-stimulated ATPase activity of *Pb*Kinesin-8B-MD, *Pb*Kinesin-8B-MD-L2^KIF5B^ and *Pb*Kinesin-8B-MD-L12^KIF5B^. Data (n=3 for each point, mean ± SD) was fitted using Michaelis-Menten equation, from which the Kcat and K_M_ were calculated in Prism9; e) MT depolymerisation rate (nm/s) for PbKinesin-8B-MD, PbKinesin-8B-MD-L2^KIF5B^ and PbKinesin-8B-MD-L12^KIF5B^ in the presence of ATP. Error bars represent the mean ± SD and individual measurements are also plotted. Ordinary one-way ANOVA was performed in Prism. Significance values are displayed as asterisks, ****, p-values <0.0001.N_PbKinesin-8B-MD_ =97 ends, N ^KIF5B^=86 ends, N ^KIF5B^=100 ends,N =85 ends.

To test the functional contributions of distinct MT contact regions of *Pb*kinesin-8B-MD, we engineered mutants in loop 2 and loop 12. These loops are shorter in kinesin-1 compared to canonical kinesin-8 (Fig. 4b, c), and we spliced the shorter loops of human kinesin-1 (KIF5B) into the *Pb*kinesin-8B-MD sequence. Both loop substitution mutants exhibited ∼40-50% of the WT ATPase activity, with Kcat-*Pb*kinesin-8B-MD-L2^KIF5B^ = 0.9 ± 0.1 ATP/s and Kcat-*Pb*kinesin-8B-MD-L12^KIF5B^ = 0.8 ± 0.3 ATP/s compared to Kcat-WT = 1.9 ± 0.2 ATP/s. Both mutants also exhibited a higher K_m_MT compared to WT (Fig. 4d). Intriguingly, while loop 2 substitution removes more amino acids, including positively charged residues that could interact with the surface of the MT, the Km of this mutant was only reduced by 2-fold. In contrast, the KmMT of the loop 12 chimera - which also has a reduced positive charge in the context of a much shorter loop – was ∼20-fold greater than WT. Presumably due to this much weaker MT interaction, the MT depolymerisation activity of *Pb*kinesin-8B-MD-L12^KIF5B^ was also much reduced compared to WT and was not significantly different from the no-kinesin control (Fig. 4e). Surprisingly, *Pb*kinesin-8B-MD-L2^KIF5B^ retained MT depolymerisation activity, albeit slower than WT. This was reinforced by the fact that, on incubation of this mutant with tubulin and AMPPNP, tubulin rings with dimensions indistinguishable from those of WT *Pb*kinesin-8B-MD were observed using negative stain EM (Supplementary Fig 7c, d). This shows that loop 2 of *Pb*kinesin-8B-MD is not required for the specific interaction and stabilisation of curved tubulin that is correlated with MT depolymerisation activity.

### The role of the kinesin neck linker in kinesin-8B-MD function

We also investigated the contribution of the kinesin-8B neck linker to motor function and compared the activities of *Pb*kinesin-8B-MD and *Pf*kinesin-8B-MD with and without (*Pb*kinesin-8B-MDΔNL and *Pf*kinesin-8B-MDΔNL) their C-terminal neck linkers (Fig. 5a). In the ATPase assay, while the K_m_ of *Pb*kinesin-8B-MDΔNL (1.8±1.0 μM) is only slightly lower than that of *Pb*kinesin-8B-MD (2.5±0.6 μM), its Kcat was substantially reduced, at 0.3 ± 0.1ATP/s compared to 1.9 ± 0.2ATP/s (Fig. 5b). Likewise, the K_m_MT and Kcat of *Pf*kinesin-8B-MDΔNL are also lower than that of *Pf*kinesin-8B-MD (K_m_MT: 0.5 ± 0.2 μM vs 1.2±0.5 μM; Kcat: 0.5±0.1ATP/s vs 1.4±0.2ATP/s (Fig. 5b). Neither *Pb*kinesin-8B-MDΔNL nor *Pf*kinesin-8B-MDΔNL generated MT gliding activity (Fig. 5c). Furthermore, the MT depolymerisation activity of both of these constructs was significantly lower than *Pb*kinesin-8B-MD and *Pf*kinesin-8B-MD (Fig. 5d). Together, these data demonstrate the importance of the kinesin-8B neck linker sequence for all of these motors’ functions.

**Figure 5.**
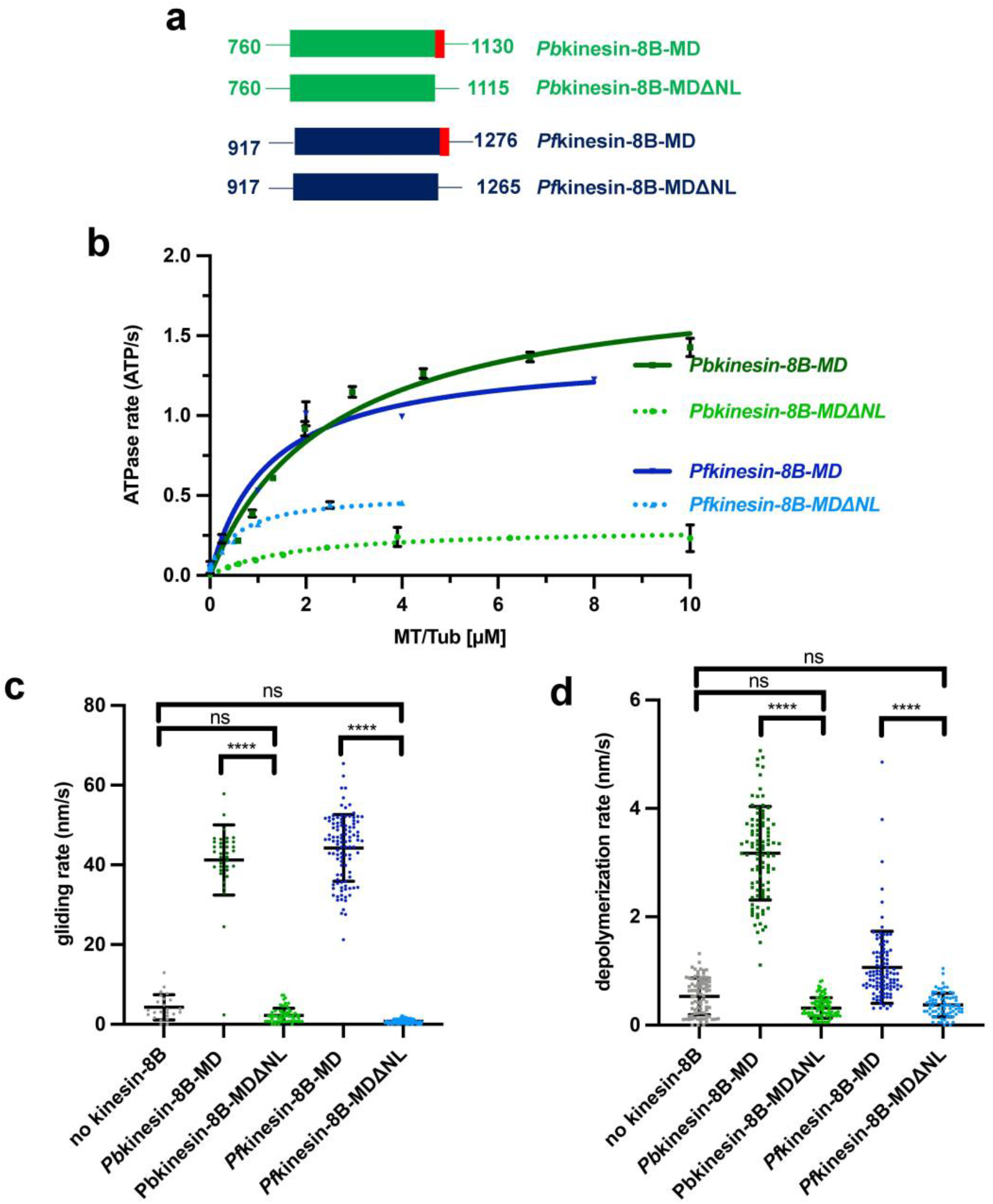
*Plasmodium* kinesin-8B neck linker is required for both motility and depolymerase activities. a) Schematic of MD and MDΔNL constructs of *Pb*kinesin-8B and *Pf*kinesin-8B. Motor domains are coloured in green and blue, respectively; neck linker sequences are coloured in red. b) Kinesin-8B-MDΔNL constructs exhibit reduced MT-stimulated ATPase activity compared to Kinesin-8B-MD. Data (n = 3 for each point) was fitted using Michaelis-Menten equation (mean +/- SD), from which the Kcat and K_M_ were calculated in Prism9. c) Kinesin-8B MDΔNL construct exhibit no significant gliding activity. Ordinary one-way ANOVA was performed in Prism. Significance values are displayed as asterisks, ****, p-values <0.0001; ***, p-values 0.0001 to 0.001. N_PbKinesin-8B-MD_ = 36 MTs. N_PbKin8-B_MDΔNL_= 67 MTs. N_PfKinesin-8B-MD_ = 104 MTs. N_PfKin8-B_MDΔNL_=77 MTs.N_no kinesin8B_= 24 MTs. d) MT depolymerisation rate (nm/s) for PbKinesin-8B-MDΔNL and PfKinesin-8B-MDΔNL in the presence of ATP compared to *Pb*kinesin-8B-MD and *Pf*kinesin-8B-MD and a no kinesin control. Error bars represent the mean ±SD and individual measurements are also plotted with coloured points. Ordinary one-way ANOVA was performed in Prism. Significance values are displayed as asterisks, ****, p-values <0.0001. N_PbKinesin-8B-MD_ =97 ends. N_PbKinesin-8B-MDΔNL_ =91 ends, N_PfKinesin-8B-MD_ =103 ends. N_PfKinesin-8B-MDΔNL_ =75 ends, N_no kinesin8B_=85 ends. Data for *Pf*kinesin-8B-MD are replotted from Fig. 2B, while data for *Pb*kinesin-8B-MD were collected in parallel with mutant activity measurement; differences in depolymerisation rates between different experiments most likely relate to different MT stability between different preps.

## DISCUSSION

Kinesin-8s are among the most widely distributed kinesin subfamilies across eukaryotes^8^, perhaps because of their functional adaptability to both move processively along MTs and to influence MT dynamics. To understand the molecular basis of *P. berghei* kinesin-8B function in parasite transmission^6,7^, we characterised its motor domain and compared it with kinesin-8B from *P. falciparum*. Our biochemical and structural data provide evidence of conserved and precisely tuned mechanochemistry in these motors, which is distinct compared to other kinesin-8s characterised to date, including those in the parasite’s mammalian hosts.

### Plasmodium kinesin-8Bs exhibit catalytic depolymerase activity

*Pb*kinesin-8B-MD and *Pf*kinesin-8B-MD behave very similarly to each other and share the ability with other kinesin-8s – including kinesin-8X from *P. berghei* and *P. falciparum*^10^ - to drive ATP-dependent plus-end directed MT gliding (Fig. 1)^11,18,19,29^. *Pb*kinesin-8B-MD and *Pf*kinesin-8B-MD both require the neck linker sequence for this activity (Fig. 5c), consistent with models of plus-end directed kinesin motility^23,24,30,31^. *Pb*kinesin-8B-MD and *Pf*kinesin-8B-MD also depolymerise stabilised MTs and, as monomeric constructs, access both MT ends via diffusion and depolymerise them. Depolymerisation by both *Pb*kinesin-8B-MD and *Pf*kinesin-8B-MD, as well as mammalian kinesin-8B (also called KIF19A^19^) is faster in the presence of ATP – i.e. it is catalytic (Fig. 2). Catalytic depolymerisation was also observed by *Plasmodium* kinesin-8Xs^10^. In contrast, *S. cerevisiae* Kip3, a kinesin-8A, MT depolymerisation is linked to suppression of motor ATPase activity^11^ and in the case of HsKIF18A_MD (another kinesin-8A), depolymerisation is more robust in the presence of the non-hydrolysable ATP analogue AMPPNP than ATP^17,18^.This suggests that a key difference in depolymerisation activities between kinesin-8As and other kinesin-8s is that kinesin-8As exhibit non-catalytic depolymerase activity. In different functional contexts, this may manifest more as modulation of MT dynamics rather than MT depolymerisation *per se*^15^. Characterisation of kinesin-8s from a range of organisms is required to solidify this distinction.

### Extended loop 2 of *Pb*kinesin-8B shows a distinct conformation when contacting the MT surface but only modestly modulates its depolymerase activity

We used cryo-EM to determine the MT-bound NN and AMPPNP structures of *Pb*kinesin-8B-MD and of NN *Pf*kinesin-8B-MD (Fig. 3). Only a few other structures of motor domains from kinesin-8s have been determined to date^11,17-19,32^, and we compared our *Plasmodium* kinesin-8B reconstructions to HsKIF18A_MD, for which most structural data are available^18^. In the NN reconstructions, the overall MT-binding footprint of both *Pb*kinesin-8B-MD and *Pf*kinesin-8B-MD are essentially indistinguishable from KIF18A_MD at the resolutions of the available reconstructions^18^. One of the distinctive features of kinesin-8s compared to other plus-end directed kinesins is an extended loop 2. In both *Pb*kinesin-8B-MD and *Pf*kinesin-8B-MD, density corresponding to loop 2 is clear, well-structured and contacts the C-terminal end of H12 of α-tubulin. Loop 2 of HsKIF18A_MD, which is 28 amino acids longer, contacts the MT surface at a similar site – however, even when contacting the MT surface, it is flexible and lacks a clearly defined structure^17,18^, and is thereby distinct from the shorter and structurally well-defined loop 2 of kinesin-8Bs visualised to date (Supplementary Fig. 7a, b)^19^. In both KIF18A^33^ and Kip3^11^, loop 2 residues are not required for MT depolymerisation activity, and contribute instead to motor processivity or MT plus-end residence time, respectively. In contrast, loop 2 residues are crucial for KIF19A depolymerase activity^19^. We show, however, that elimination of the *Pb*kinesin-8B-MD loop 2 sequence reduces MT affinity and depolymerase activity but does not eliminate them. In summary, while the extended nature of kinesin-8 loop2 sequences likely contributes to their phylogenetic co-classification and can form an additional contact point with the MT surface, this region differently modulates motor function in different kinesin-8s.

### *Pb*kinesin-8B-MD undergoes a structurally minimal response upon AMPPNP binding

A further striking difference between MT-bound *Pb*kinesin-8B-MD and HsKIF18A_MD is its structurally minimal response to AMPPNP binding (Fig. 3i). In contrast, AMPPNP binding to HsKIF18A_MD induces rearrangements within the motor domain that support neck linker docking towards the MT plus end (maximum RMSD = 17.4^18^). The minimal structural response of *Pb*kinesin-8B-MD is surprising given the neck linker dependence of MT gliding activity by this motor (Fig. 5c) and its relatively conserved neck linker sequence (Supplementary Fig.8). It is possible that the observed small shift in the P-loop domain on AMPPNP binding (Fig. 3i) is sufficient to bias the neck linker towards the MT plus end and thereby support ATP-driven MT gliding (Fig. 1c). However, the characteristic hydrolysis-competent ‘closed’ NBS conformation^34^ isn’t observed in our AMPPNP reconstruction; it is thus also possible that AMPPNP as an analogue does not induce motility-relevant conformational changes in MT lattice bound *Pb*kinesin-8B-MD. AMPPNP binding is, however, sufficient to stabilise tubulin in a curved conformation and induce depolymerisation at MT ends (Fig. 2). A recent study of the Kip3 truncated motor domain (no neck linker) reported similar overall observations^32^ – AMPPNP binding did not induce canonical conformational changes in lattice-bound Kip3, but molecular dynamics simulations indicated that such an ATP-dependent canonical conformational change would occur at MT ends. However, a crucial difference in the behaviours observed by these motors is that AMPPNP binding to Kip3 does not induce MT depolymerisation, further emphasising that their precise mechanochemistry is distinct^11,32^.

### *Plasmodium* kinesin-8Bs share some structural features with kinesin-13s

Intriguingly, a minimal structural response to AMPPNP binding by lattice-bound kinesin-13s has also been observed^35^, but AMPPNP binding does induce depolymerisation at MT ends. Kinesin-13s are well-conserved MT catastrophe factors with regulatory roles in both interphase and dividing cells^36^. In contrast to kinesin-8s, however, kinesin-13s do not take steps along the MT lattice but diffuse to either MT end to stimulate MT depolymerisation, an activity that depends on the motor ATPase^37^. In addition – and as is the case for *Pb*kinesin-8B-MD, *Pf*kinesin-8B-MD and KIF19A – kinesin-13s are catalytic depolymerases ^20,35,38,39^. Despite the mechanistic differences with respect to lattice-based stepping, we hypothesised that the minimal response by lattice-bound kinesin-13s and *Pb*kinesin-8B-MD to AMPPNP binding could reflect a distinct mechanochemical sensitivity of catalytic depolymerases to the underlying tubulin substrate. Intriguingly, when motor domain structures of NN MT-bound *Pb*kinesin-8B-MD, HsKIF18A_MD and the MT-bound *Drosophila melanogaster* kinesin-13 KLP10A_MD (*Dm*KLP10A_MD) are overlaid by alignment on their tubulin-binding subdomains (Fig 6a), the position of the rest of *Pb*kinesin-8B-MD (by relative angle between α-helices and helix-α4) is more similar to *Dm*KLP10A_MD than to HsKIF18A_MD (Fig. 6b). Thus, while at the primary sequence level, kinesin-8s are more similar to each other (as expected from their family classification – Supplementary Fig.8), the structural comparison of motor domains, suggests that the configuration of *Plasmodium* kinesin-8Bs – and we speculate kinesin-8Bs more generally – shares some features with kinesin-13s and specifies their mechanochemistry (Fig. 6c).

**Figure 6.**
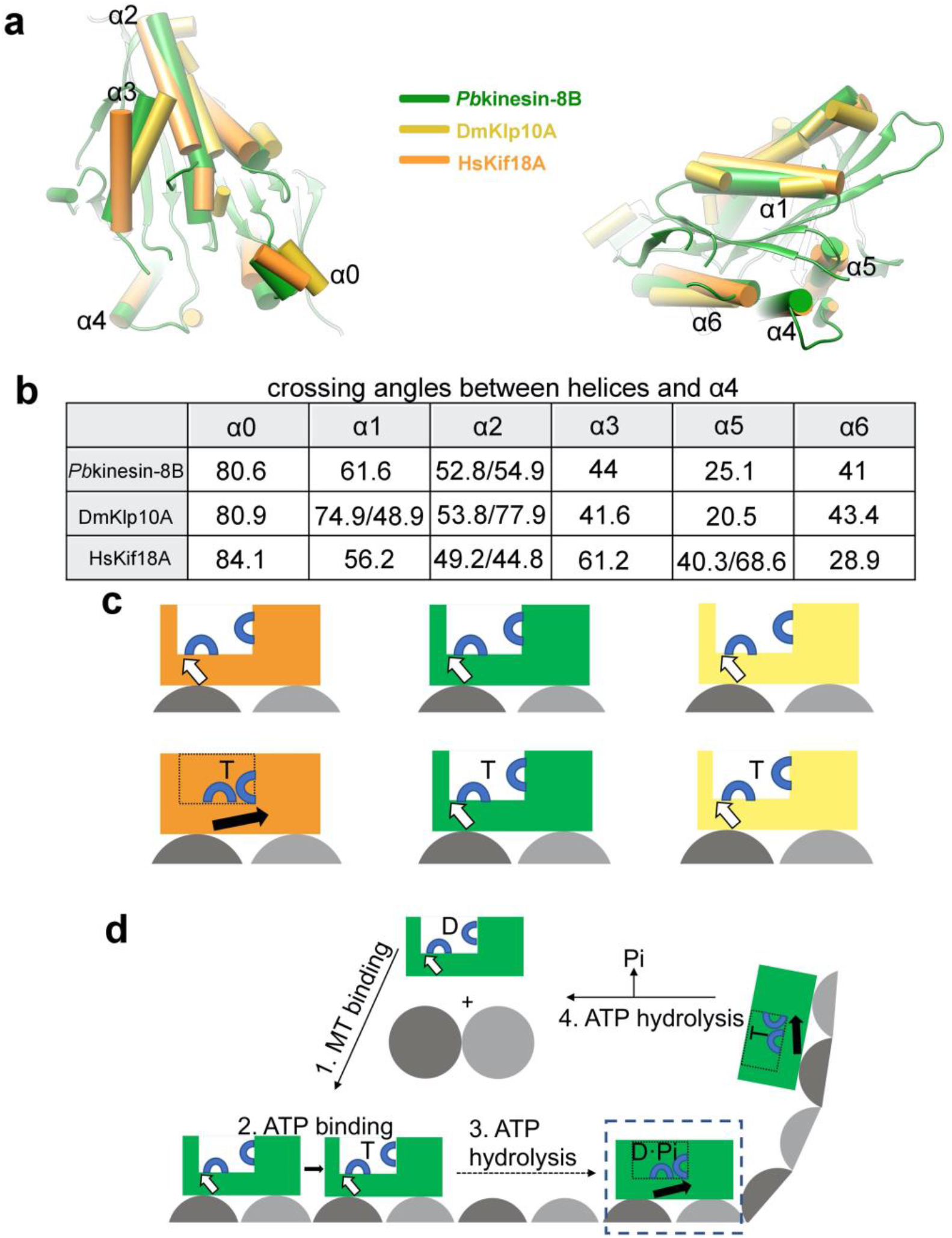
Structural comparison of MT-bound *Plasmodium* kinesin-8B with kinesin-8A and kinesin-13 motor domains. a) Structural alignment of MT-bound NN *Pb*kinesin-8B-MD with *Hs*KIF18A_MD (kinesin-8A) and *Dm*KLP10A_MD (kinesin-13) viewed towards the NBS (left) and towards the neck linker (right); b) Comparison of angle between motor domain α-helices and helix-α4 in each motor domain; c) Schematic comparison of lattice-bound motor domain responses to ATP binding in HsKIF18A (orange), Pbkinesin-8B (green), KIF18A (orange), KLP10A (yellow); d) Schematic of *Plasmodium* kinesin-8B (green) nucleotide-dependent motile and depolymerase activities 1) ADP-bound motor binds MT lattice and releases ADP (analogous to other kinesins^75^); 2) ATP binding does not induce large global conformational changes in lattice-bound motor; 3) conformational changes at other points in the ATPase cycle (e.g. ATP hydrolysis depicted here^75^) supports neck linker docking and thereby motility towards the MT plus end; 4) At MT plus ends, ATP binding induces tubulin release, ATPase turnover causes motor release from tubulin and motors are thus recycled for further activity. The MT-bound step we hypothesise exists but didn’t structurally characterise is boxed.

Taken together, these data suggest that the motor activity of *Plasmodium* kinesin-8Bs has evolved to be both motile and capable of catalytic MT depolymerisation (Fig. 6d). ATPase-dependent conformational changes in lattice-bound motors – probably not all of which were captured in our current study – bias motor movement towards the MT plus-end (Fig. 6d, step 1-3). At the MT end, larger conformational changes are enabled and drive catalytic ATP-dependent depolymerisation (Fig. 6d, step 4). In parasites, the context in which the kinesin-8B motor domain operates is likely to influence this finely tuned activity – in the context of the full-length motor, when interacting with *Plasmodium* tubulin^40^ and in the cellular environment, potentially modulated by binding partners and cellular regulators^41^.

### A conserved role for catalytic depolymerases in male gamete formation

*Plasmodium* kinesin-8Bs are expressed exclusively in male gametocytes, the only flagellated stage of the parasite life cycle. While deletion of the *P. berghei* kinesin-8B has no effect on parasite blood stages^42^ and also has no effect on genome duplication during male gametogenesis, kinesin-8B knockout parasites exhibit disrupted flagella assembly and parasite transmission is blocked^6,7^. The motor domains from *P. berghei* and *P. falciparum* kinesin-8Bs exhibit very similar properties *in vitro* which, together with their very similar expression profiles^43^, suggest a conserved function for kinesin-8Bs across *Plasmodium* species. In contrast to the well-described mechanisms of cilia and flagella assembly and maintenance that involve intraflagellar transport (IFT), the axonemes of *Plasmodium* male gametes are assembled within the cytoplasm; this process is thought to be a simplification of sperm assembly mechanisms seen in other organisms^44^. In *P. berghei* kinesin-8B knockout parasites, axoneme MT doublets accumulate in the male gamete cytoplasm but never combine to form a mature flagella^7^. This observation, together with kinesin-8B’s localisation to basal bodies embedded in the nuclear membrane at the initiation of male gametogenesis^45^, has suggested that this motor is involved in basal body maturation prior to axoneme assembly^7^. While the detailed mechanisms involved in male gamete axoneme assembly are lacking, it is striking that the kinesin-13 *Dm*KLP10A also plays a distinct role in sperm development and associates with basal bodies in the early stages of spermatogenesis. Axoneme elongation is disrupted when KLP10A expression is reduced^46^, hinting at a conserved role for regulation of MT length or dynamics in basal body regulation early in male gamete formation. Subsequent association of *P. berghei* kinesin-8B along the length of the mature axoneme suggests that this motor has additional roles in the mature flagella of male gametes.

Our characterisation of kinesin-8B motor domains from *P. berghei* and *P. falciparum* emphasises the wider importance of MT length regulation at all stages of axoneme development and function in cilia/flagella. Several kinesin families, including kinesin-8s and kinesin-13s in diverse organisms^47-52^ have been implicated in these processes. Our study highlights the utility of kinesin group classifications as a starting point for investigation of the molecular mechanism of motors that regulate MT organelle size, but also the fluidity of those molecular properties in particular functional contexts. Future studies will reveal the cellular context in which these enzymes are utilised, leading to better understanding of divergence from canonical mammalian kinesins. Molecular understanding of these motors can thereby also contribute to potential new inhibition strategies for blocking key parasite life cycle transitions that rely on kinesins^53^.

## MATERIALS AND METHODS

### Molecular cloning

DNA encoding the motor domain of *Pb*kinesin-8B (PBANKA_020270), residues 760-1130, referred to as *Pb*kinesin-8B-MD, was codon optimized, synthesized (Gene Oracle, Inc.) and cloned into the pNIC28-Bsa4 vector (Structural Genomics Consortium, Oxford, UK) using ligation independent cloning. *Pb*kinesin-8B-MDΔNL (residues 760-1115) was prepared by introduction of a stop codon using site-directed mutagenesis after the codon for residue 1115. For the *Pb*kinesin-8B-MD-SNAP construct and *Pb*kinesin-8B-MDΔNL-SNAP construct, Gibson assembly was used to insert a SNAP_f_-tag at the C-termini of these constructs. A flexible linker of (GGGS)2 was added immediately before the C-terminal SNAP_f_-tag of *Pb*kinesin-8B-MDΔNL-SNAP construct. The *Pb*kinesin-8B-MD ^E1023A^ was generated using site-directed mutagenesis. The *Pb*kinesin-8B-MD_L2^KIF5B^ construct was generated by replacing the loop2 sequence (LDPSDNTDNVLRQNRTKE) in *Pb*kinesin-8B-MD with the corresponding sequence (AS) from human KIF5B using Gibson assembly. Similarly, the *Pb*kinesin-8B-MD_L12^KIF5B^ construct was generated by replacing the loop12 sequence (SRSKGTSKSNFIPF) of *Pb*kinesin-8B-MD with the corresponding sequence (EGSTYVPY) from human KIF5B using Gibson assembly.

DNA encoding the motor core of *Pf*kinesin-8B (PF3D7_0111000), residues 917-1265, referred to as *Pf*kinesin-8B-MDΔNL, was codon optimized, synthesized (Gene Oracle, Inc.) and cloned into the pNIC28-Bsa4 vector using ligation independent cloning. The putative neck linker sequence, residues 1266-1276, was incorporated into this construct using Gibson assembly and referred to as *Pf*kinesin-8B-MD. Insertion of a SNAP_f_-tag at the C-terminus of the constructs was also performed using Gibson cloning. Due to poor solubility of *Pf*kinesin-8B-MD_SNAP_f_ on expression in *E. coli*, an N-terminal NusA solubility tag ^54^ was also incorporated into this construct, C-terminal of the 6xHis tag but N-terminal of the TEV protease cleavage site so that it could be removed during purification.

### Protein expression

For expression of all *Pb*Kinesin-8B proteins, expression plasmids were transformed into BL21 Star™ DE3 *E. coli* competent cells (Invitrogen). Cells were grown to OD 0.6-0.8 at 37°C then induced with 20 μM IPTG. After induction, the temperature was lowered to 20°C and cells were incubated overnight before pelleting by centrifugation at 6,235 g for 20 mins.

For expression of *Pf*kinesin-8B-MD and *Pf*kinesin-8B-MDΔNL proteins, expression plasmids were transformed into BL21 Star™ DE3 *E. coli* competent cells (Invitrogen). Cells were grown to OD 0.6-0.8 at 37°C then induced with 1mM IPTG. After induction the temperature was lowered to 26°C and cells were incubated overnight before pelleting by centrifugation at 6,235g for 20mins. For expression of *Pf*kinesin-8B-MDΔNL_SNAP and NusA_*Pf*kinesin-8B-MD_SNAP, cells were induced with 100uM IPTG induction and were grown at 18°C or 26°C respectively post-induction.

### Protein purification

For purification of all *Pb*kinesin-8B related proteins, pelleted cells were resuspended in lysis buffer (20 mM Tris-HCl pH 7.5, 500 mM NaCl, 5 mM MgCl_2_ with EDTA-free protease inhibitor (Roche)), sonicated for 30 min and then centrifuged at 48,384 g, 4°C for 30 min. The His_6_- tagged proteins were purified using Immobilised Metal Affinity Chromatography (IMAC) with 2mL Ni-NTA His•Bind^®^ Resin (Merck), followed by incubation with TEV protease for 12 hours at 4 °C to remove the His_6_ tag. The protein was exchanged into low-salt buffer (20 mM Tris-HCl pH 7.5, 100 mM NaCl, 5 mM MgCl2) and subjected to a further reverse-IMAC step, followed by application to HiTrap Q HP IEX column (GE Healthcare) to remove any residual bacterial proteins, which bind to the Q column. The Q column flow-through was collected and concentrated using Amicon Ultra-0.5ml Centrifugal Filters (Millipore) then separated into single-use aliquots, snap frozen in liquid nitrogen and stored at -80°C.

For purification of *Pf*kinesin-8B proteins, pelleted cells were resuspended in lysis buffer (50mM Tris pH7.0, 400mM NaCl, 2mM MgCl_2_, 1mM ATP, 2mM beta-mercaptoethanol, 15µg/ml DNase I (Sigma) with EDTA-free protease inhibitor (Roche)), lysed by passage 3 times through an Avesti Emusiflex C3 high-pressure homogeniser and centrifuged at 48,384 g for 1 hour. The His_6_-tagged proteins were purified using Immobilised Metal Affinity Chromatography (IMAC) with Ni-NTA His•Bind^®^ Resin (Merck). Fractions containing the protein of interest were then dialysed against low-salt buffer (50mM Tris pH7.0, 40mM NaCl, 2mM MgCl_2_, 1mM ATP, 2mM beta-mercaptoethanol) for 12 hours at 4°C. TEV protease was added during dialysis to remove the N-terminal His_6_-tag.The protein was retrieved from dialysis and loaded onto a 1ml HiTrap SP HP IEX column (GE Healthcare) and eluted by gradient introduction of high-salt buffer (50mM Tris pH7.0, 1M NaCl, 2mM MgCl_2_, 1mM ATP, 2mM beta-mercaptoethanol) on an ÅKTA system (GE Healthcare). Protein-containing pooled fractions from IEX were then loaded onto a Superdex 200 Increase 10/300 GL gel filtration column (GE Healthcare) and collected in gel filtration buffer (20mM PIPES pH6.8, 80mM KCl, 2mM MgCl_2_, 1mM ATP, 2mM beta-mercaptoethanol). Fractions of monomeric protein were collected and concentrated to around 30-50 µM using Amicon Ultra-0.5 ml Centrifugal Filters (Millipore), then separated into single-use aliquots, snap frozen in liquid nitrogen and stored at -80°C.

In the specific case of NusA_*Pf*kinesin-8B-MD_SNAP, the pI of the protein was close to pH7.0 therefore precipitation was observed with the purification method above. Instead pH8.5 was used for all purification buffers. Pelleted cells were resuspended in lysis buffer (50mM Tris pH8.5, 400mM NaCl, 2mM MgCl_2_, 1mM ATP, 2mM beta-mercaptoethanol, 15µg/ml DNase I (Sigma) with EDTA-free protease inhibitor (Roche)). Resuspended cells were lysed using high-pressure homogeniser, followed by centrifugation at 48,384 g for 1 hour as above. The His_6_-tagged proteins were purified using Immobilised Metal Affinity Chromatography (IMAC) with Ni-NTA His•Bind^®^ Resin (Merck). Fractions containing the protein of interest were then dialysed against low-salt (50mM Tris pH8.5, 40mM NaCl, 2mM MgCl_2_, 1mM ATP, 2mM beta-mercaptoethanol) together with TEV protease treatment to cut both His_6_ and NusA tags. The *Pf*kinesin-8B-MD construct retrieved from dialysis was loaded onto a 1ml HiTrap 1ml Q FF IEX column (GE healthcare). *Pf*kinesin-8B-MD_SNAP was eluted by gradient introduction of high-salt buffer (50mM Tris pH8.5, 1M NaCl, 2mM MgCl_2_, 1mM ATP, 2mM beta-mercaptoethanol). Protein-containing pooled fractions from IEX were then loaded onto a Superdex 200 Increase 10/300 GL gel filtration column (GE Healthcare) and collected in gel filtration buffer (20mM PIPES pH6.8, 80mM KCl, 2mM MgCl_2_, 1mM ATP, 2mM beta-mercaptoethanol). Fractions containing the protein of interest were collected and concentrated to around 30-50µM using Amicon Ultra-0.5ml Centrifugal Filters (Millipore) then separated into single-use aliquots, snap frozen in liquid nitrogen and stored at -80°C.

### MT polymerization

For all assays, porcine brain tubulin was purchased as a lyophilised powder (Cytoskeleton, Inc.) either unlabelled, X-rhodamine-labelled or biotinylated. The protein was solubilized in BRB80 buffer (80 mM PIPES-KOH pH 6.8, 1 mM EGTA, 1 mM MgCl_2_) to approximately 10 mg/ml (tubulin dimer concentration).

#### Paclitaxel-stabilized MTs

Reconstituted tubulin was polymerised at 5 mg/ml final concentration in the presence of 5 mM GTP at 37 °C for 1 hour. After this, a final concentration of 1 mM paclitaxel (Calbiochem) dissolved in DMSO was added and the MTs incubated at 37 °C for a further 1 hour.

#### GMPCPP-stabilised MTs

GMPCPP MTs were prepared using a double-cycling protocol as follows to maximise GMPCPP occupancy. Reconstituted tubulin was polymerised at 5 mg/ml final concentration in the presence of 1 mM GMPCPP at 37 °C for 1 hour. Polymerised MTs were pelleted at 313,000g for 10 min at 25°C using TLA100 rotor (Beckman Coulter), and the pellet was washed with BRB80 buffer. The MT pellet was then resuspended with BRB80 buffer, followed by Incubation on ice for 20 min to depolymerize the MTs. The mix were incubated on ice for another 5 min with an additional 1mM GMPCPP. The reaction mix was then incubated at 37°C for 30 min.

For ATPase assays, Paclitaxel-or GMPCPP-stabilized MTs were polymerized as above, and free tubulin was removed by pelleting the MTs by centrifugation at 313,000g for 10min at 25°C using TLA100 rotor (Beckman Coulter) through a sucrose cushion, the supernatant removed and the MT pellet was resuspended in BRB80 buffer. Protein concentration was determined using a Bradford assay.

For depolymerization assays, paclitaxel-stabilised MTs containing 10% X-rhodamine-labelled and 10% biotin-labelled tubulin (Cytoskeleton) were polymerized as above and were left at room temperature for 48 hours before use in the TIRF assay.

Paclitaxel-stabilised MTs and GMPCPP-polarized MTs were used in the gliding assay. Paclitaxel MTs containing 10% X-rhodamine-labelled tubulin were polymerized as above and left for 48 hours at room temperature before use in a TIRF assay. To prepare polarized MTs to detect gliding directionality, long “dim” MTs were first polymerized by mixing X-rhodamine-labelled tubulin and unlabelled tubulin at a 1:9 ratio to a final concentration of 2 mg/ml. This mix was incubated at 37 °C for 2 hours in the presence of 0.5 mM GMPCPP. MTs were then pelleted by centrifugation at 17,000g in a bench-top centrifuge for 15 min. To add bright plus end caps to the MTs, X-rhodamine-labelled tubulin and unlabelled tubulin were mixed in a 1:1 ratio. The unlabelled tubulin in this reaction had been previously incubated with 1 mM N-ethyl maleimide (NEM) on ice for 10 min, followed by incubation with 8 mM beta-mercaptoethanol on ice for 10 min to block growth from the MT minus-end. This “bright” mix was pre-warmed then added to the polymerized long, dim MTs and incubated at 37 °C for 15 min. MTs were pelleted by centrifugation and resuspended in BRB80 with 40 μM paclitaxel.

### MT-and tubulin-stimulated ATPase assay

MT/tubulin-stimulated kinesin ATPase activity was measured using an NADH-coupled assay^55^. The assay was performed using 250 nM *Pb*kinesin-8B-MD or *Pf*kinesin-8B-MD titrated with paclitaxel-stabilised MTs (*Pf*), GMPCPP MTs (*Pb*) or tubulin dimer(*Pb*) in 100 μl ATPase reaction buffer containing an ATP regeneration system: For *Pb*Kinesin-8B-MD: BRB80 buffer, 5mM ATP(Sigma), 5 mM phosphoenolpyruvate (PEP), 2 mM NADH, 12 U pyruvate kinase and 16.8 U lactate dehydrogenase; for *Pf*Kinesin-8B-MD: BRB80 buffer, 5mM ATP(Sigma), 50mM NaCl, 5 mM phosphoenolpyruvate (PEP), 2 mM NADH, 12 U pyruvate kinase and 16.8 U lactate dehydrogenase. NADH depletion was monitored by the decrease in absorbance at 340 nm in a SpectraMax Plus-384 plate reader at 26 °C operated by SoftMax Pro 5 software. The Michaelis-Menten equation was used for curve fitting of the ATPase data using Prism 9. To compare ATPase rates between *Pb*kinesin-8B-MD and its mutants, all *Pb*kinesin-8B proteins were buffer exchanged to BRB80 buffer before use in the ATPase assay and the assay was performed in BRB80 buffer containing the above ATP regeneration system without addition of any NaCl.

### MT gliding assay

SNAP–tagged kinesin-8B-MD proteins (20 μM) were biotinylated in 50 μl reaction volumes by incubating with 40 μM SNAP-biotin (NEB) at 4 °C overnight. Proteins were purified from excess SNAP-biotin by size-exclusion chromatography on a Superdex 75 Increase 3.2/300 column using an ÅKTA micro system (GE Healthcare) in gel filtration buffer (20 mM Tris-HCl pH 7.5, 250 mM NaCl, 5 mM MgCl_2_, 1 mM DTT). Peak fractions were pooled, snap frozen in liquid nitrogen and stored at -80°C.

Flow chambers for Total Internal Reflection Fluorescence (TIRF) microscopy were made between glass slides, biotin-PEG coverslips (MicroSurfaces Inc.), and double-sided tape. Chambers were sequentially incubated with: 1) blocking solution (0.75 % Pluronic F-127, 5 mg/ml casein) for 5 min, followed by two washes with assay buffer (BRB80 buffer, 1 mM DTT and 20 μM paclitaxel); 2) 0.5 mg/ml neutravidin for 2 min, followed by two washes with assay buffer (BRB80 buffer, 1 mM DTT and 20 μM paclitaxel); 3) biotinylated kinesin-8B-MD, incubated for 2 min, followed by two washes with assay buffer supplemented with 1 mg/ml casein; 4) the reaction mixture containing 5mM ATP together with 10 % X-rhodamine-MTs (or polarity marked GMPCPP MTs to determine directionality) in assay buffer supplemented with an oxygen scavenging system (20 mM glucose, 300 μg/ml glucose oxidase, 60 μg/ml catalase).

An Eclipse Ti-E inverted microscope was used with a CFI Apo TIRF 1.49 N.A. oil objective, Perfect Focus System, H-TIRF module, LU-N4 laser unit (Nikon) and a quad band filter set (Chroma)^56^. Movies were collected at 26°C under illumination at 561 nm for 10 min with a frame taken every 2 s with 100 ms exposure on a iXon DU888 Ultra EMCCD camera (Andor), using the NIS-Elements AR Software (Nikon). The gliding rates of single MTs were measured from kymographs using Fiji software^57^.

### MT depolymerisation assay

Flow chambers were treated and incubated with blocking solution and washed twice with assay buffer (BRB80 buffer, 1 mM DTT and 20 μM paclitaxel), followed by incubation with 0.5 mg/ml neutravidin and two washes with assay buffer as above in MT gliding assay. 1:100 dilution of X-rhodamine and biotin-labelled paclitaxel-stabilized MTs were flooded into the chamber and incubated for 2 min, followed by two washes with assay buffer supplemented with 1 mg/ml casein; 5 μM unlabelled kinesin-8B-MD and mutants in assay buffer supplemented with 5 mM nucleotide (as indicated) and an oxygen scavenging system (20 mM glucose, 300 μg/ml glucose oxidase, 60 μg/ml catalase) were introduced into chamber right before observation. For assays comparing *Pb*kinesin-8B mutants, all *Pb*kinesin-8B related proteins were buffer exchanged into BRB80 buffer prior to use in the assay. The microscope and camera used were the same as for the MT gliding assays. Movies were collected at 26°C under illumination at 561 nm for 30 min with a frame taken every 10 s with 100 ms exposure. MT depolymerisation rates were determined from kymographs using Fiji software.

### Negative stain sample preparation, data collection and analysis of tubulin ring structures

60 μM Pbkinesin-8B-MD or Pbkinesin-8B-MD-L2^KIF5B^ were incubated with 20 uM tubulin in BRB80 buffer in the presence of 5mM AMPPNP at room temperature for 1h. The reaction mix was diluted into BRB80 buffer 10-fold, followed by application of 4 μl of the mix onto glow discharged continuous carbon electron microscopy grid (400 mesh, EMS) and was incubated on grid for 1 min. The sample drop was blotted using filter paper (Whatman) before 4 μl of 2% uranyl acetate was applied. After incubation on grid for a further 1min, the stain was blotted with filter paper and the grid was allowed to dry. All negative stain micrographs were collected using a Tecnai T12 transmission electron microscope (Thermo Fisher Scientific) with a 4×4K CCD camera (Gatan) at 120 kV, using magnification of 52,000, with an image pixel size of 2.09 Å and defocus around -5 μm. Diameters of tubulin rings were measured in Fiji^57^.

### Cryo-EM sample preparation

25 μM *Pb*kinesin-8B-MD or 50 μM *Pf*kinesin-8B-MD was incubated in BRB80 buffer containing apyrase (10 units/ml, P*b* and P*f*) or 5mM AMPPNP (P*b*) at room temperature for 15min. 4 μl of 10 μM GMPCPP-MT (polymerised as described above) were applied to a glow-discharged C-flat 2/2-4C grid (EMS) at room temperature. After incubation on the grid for 1 min, 3.5ul of MTs were removed by pipetting, followed by double application of 3.5ul of the *Pb*kinesin-8B-MD mix. Grids were then plunge frozen using Vitrobot Mark IV (Thermo Fisher Scientific) with the following setting: blot force of 5, blot time of 5s, humidity of 100%, and temperature of 22°C.

For the tubulin ring samples, 60 μM *Pb*kinesin-8B-MD was incubated with 20 μM tubulin and 5mM AMPPNP at room temperature for 1hour. 4 μl were applied to a glow-discharged C-flat 2/2-4C grid. Grids were then plunge frozen using Vitrobot Mark IV (Thermo Fisher Scientific) with the following setting: blot force of 5, blot time of 5s, wait time 10s, humidity of 100%, and temperature of 22°C.

### Cryo-EM data acquisition

For the *Pb*kinesin-8B-MD NN dataset, 337 movies were collected on a Tecnai G2 Polara microscope (Thermo Fisher Scientific) with K2 Summit detector operating in counting mode with a GIF Quantum LS Imaging Filter (Gatan). The microscope was operated at an accelerating voltage of 300 kV with nominal magnification of 160K and pixel size of 1.35Å. 50 frames for each micrograph were collected using serialEM software^58^, 15s exposure time, 51 e^-^/ Å^2^ total electron exposure dose and 7 e-/pixel/s dose rate. The defocus range is from -0.5 to -2.5 μm.

For *Pb*kinesin-8B-MD AMPPNP dataset, 1026 movies were collected on a Titan Krios microscope (Thermo Fisher Scientific) with K2 Summit detector operating in counting mode with a GIF Quantum LS Imaging Filter (Gatan). The microscope was operated at an accelerating voltage of 300 kV with nominal magnification of 130K and pixel size of 1.05Å. 32 frames for each micrograph were collected using EPU (Thermo Fisher Scientific), 8s exposure time, 47e^-^/ Å^2^ total electron exposure dose and 7e-/pixel/s dose rate. The defocus range is from -0.5 to -2.5 μm. The *Pf*kinesin-8B-MD NN dataset consisted of 4075 movies which were collected similarly, using 8s exposure time, 47 e^-^/ Å^2^ total electron exposure dose and 7e^-^/pixel/s dose rate. The defocus range is from -0.5 to -2.5 μm.

For the *Pb*kinesin-8B-MD-tubulin cryo-EM dataset, 8148 movies were collected on a Titan Krios microscope (Thermo Fisher Scientific) with K2 Summit detector operating in counting mode with a GIF Quantum LS Imaging Filter (Gatan). The microscope was operated at an accelerating voltage of 300 kV with nominal magnification of 105K and pixel size of 1.37 Å. 40 frames for each micrograph were collected with using EPU (Thermo Fisher Scientific), 12s exposure time, 40 e^-^/ Å^2^ total electron exposure dose and 6e-/pixel/s dose rate. The defocus range is from -0.5 to -2.5 μm.

### Data processing

Movie frames were motion-corrected using MotionCor2 as follows: *Pb*kinesin-8B-MD NN: frame 2-24, *Pb*kinesin-8B-MD AMPPNP: frame 2-16, *Pf*kinesin-8B-MD NN: frame 1-32. The parameters of the contrast transfer function (CTF) for each micrograph were determined using CTFFIND4 ^59^. Particles with box size of 432 pixels (*Pb*kinesin-8B-MD NN), 576 pixels (*Pb*kinesin-8B-MD AMPPNP, *Pf*kinesin-8B-MD NN) and non-overlapping region of one tubulin dimer size were picked using EMAN2 e2helixboxer.py ^60^. All picked particles were imported into RELION v3.0 and performed using the MiRP pipeline ^61-63^. Briefly, 4x binned particles were subjected to supervised 3D classification with 15 Å low-pass filtered references of MTs with different protofilament numbers. Particles from each MT were then assigned a unified protofilament number class and 14 protofilament MTs were taken for further processing. Next, several rounds of 3D alignment followed by smoothing of Euler angles and X/Y shifts assignments based on the prior knowledge of MT architecture was performed. From this, an initial seam location was determined for each MT, which was then checked and corrected using supervised 3D classification against references for all possible seam locations. C1 reconstruction was performed on un-binned particles using auto-refine with alignment parameters obtained from above processing steps (Table 1).

Low occupancy and/or flexibility of the motor domains on the MT lattice results in a resolution decay in the final reconstructions from MT to kinesin density. To improve the density of kinesin, further steps were performed as previously described^25^. First, symmetry expansion was performed in RELION with these unbinned particles to obtain the 14-fold expanded dataset. After this step, only the protofilament opposite the MT seam exhibits the correct αβ-tubulin registration. Density of the central kinesin-bound tubulin dimer on this protofilament was subjected to focused 3D classification without alignment (4 classes, T=256), which resulted in classes with good kinesin density and classes with no/poor kinesin density. The particles in classes without kinesin density or with poor kinesin density were discarded. The remaining particles were used for final round of 3D auto-refine on the entire density. All reconstructions were sharpened and filtered using RELION Local resolution. Density of the central kinesin-bound tubulin dimer on the protofilament opposite the MT seam was used for model building and interpretation.

For 2D analysis of the *Pb*kinesin-8B-MD-tubulin ring cryo-EM data, processing was performed in Cryosparc^64^. Frames were motion-corrected using patch motion, followed by CTF estimation using CTFFIND4. Particles were initially selected manually for 2D classification. The best classes were selected as templates for template picking. 89,836 particles were picked and extracted for multiple rounds of 2D classification.

### Model building

100 comparative models of *Pb*Kinesin8B-MD were calculated using MODELLER v9.23 ^65^, using multiple known structures as templates (PDB IDs: 5GSZ^19^, 4LNU ^30^, 3HQD^34^, and 4OZQ^66^. The top 10 models were selected using SOAP scoring ^67^, then the top model selected using QMEAN ^68^. The comparative models were rigidly fitted into no nucleotide and AMPPNP reconstructions using the *Fit-in-Map* tool in Chimera ^69^. To improve the fit to the density, a local all-atom fit to density step was performed using Rosetta Relax, incorporating a fit to density term^70^. To improve models of loop2 (16 Amino Acids(AAs)), loop5 (6 AAs), loop11 (13 AAs), the visible neck linker (5 AAs), and the N-terminus (5 AAs), loop conformations were predicted using Rosetta. First, 500 models using cyclic coordinate descent with fragment insertion were calculated ^71^, then the model with highest cross correlation to the cryo-EM density was selected. From this top model a further 500 models were calculated using kinematic closure with a fit-to-density term and the top model selected based on cross-correlation ^72^.

A *Pf*Kinesin8B-MD-NN model was generated by mutating amino acids using the “mutate” tool in Coot from *Pb*Kinesin8B-MD-NN model, which exhibits 88% sequence identity and 94% similarity^73^ (Supplementary Fig. 1). Cross correlation with the cryo-EM density was calculated showing a good fit, while the calculated QMEAN value and Molprobity score demonstrated the good geometry of the model (Table 2).

### Sequence alignment

Sequence alignments were performed with Clustal Omega, with residue colouring according to the Clustal X scheme^74^.

## DATA AVAILABILITY

The MT-bound *Pb*kinesin-8B-MD_NN, *Pb*kinesin-8B-MD_AMPPNP and *Pf*kinesin-8B-MD_NN reconstructions will be deposited with the Electron Microscopy Data Bank, deposition number *****, ***** and ***** respectively. The molecular models of MT-bound *Pb*kinesin-8B-MD_NN, *Pb*kinesin-8B-MD_AMPPNP and *Pf*kinesin-8B-MD_NN will be deposited with the Worldwide Protein Data Bank, deposition number *****, ***** and ***** respectively.

## ACKNOWLEDGEMENTS

This work was supported by grants from the Biotechnology and Biological Sciences Research Council, U.K. (BB/N018176/1 to C.A.M.) and the Wellcome Trust (101311-10 to A.D.C.; 104196/Z/14/Z and 217186/Z/19/Z to A.J.R; 085945/Z/08/Z) to C.A.M.). F.S. and A.D.C. were supported by a PhD studentship from the Biotechnology and Biological Sciences Research Council, U.K. and Medical Research Council, U.K respectively. R.T. was supported by the Biotechnology and Biological Sciences Research Council, U.K. (BB/N017609/1). C.J.S. is supported by the UK Health Security Agency, the EDCTP WANECAM II Consortium and the Medical Research Council, U.K. (MR/T016124.1). Cryo-EM data collected at the Institute of Structural and Molecular Biology (ISMB), Birkbeck was on equipment funded by the Wellcome Trust, U.K. (202679/Z/16/Z, 206166/Z/17/Z and 079605/Z/06/Z) and the Biotechnology and Biological Sciences Research Council (BBSRC) UK (BB/L014211/1). We thank N. Lukoyanova and S. Chen for electron microscope support, D. Houldershaw for computing support at Birkbeck, M. Topf for advice on structural modelling, and members of the Moores group for helpful discussions

## COMPETING INTERESTS

Fiona Shilliday declares that she is now an employee of AstraZeneca. The other authors declare no competing interests

## REFERENCES

1 Gerald, N., Mahajan, B. & Kumar, S. Mitosis in the human malaria parasite Plasmodium falciparum. Eukaryot Cell 10, 474–482, doi:10.1128/EC.00314-10 (2011).

2 Frénal, K., Dubremetz, J. F., Lebrun, M. & Soldati-Favre, D. Gliding motility powers invasion and egress in Apicomplexa. Nat Rev Microbiol 15, 645–660, doi:10.1038/nrmicro.2017.86 (2017).

3 Sinden, R. E., Talman, A., Marques, S. R., Wass, M. N. & Sternberg, M. J. The flagellum in malarial parasites. Curr Opin Microbiol 13, 491–500, doi:10.1016/j.mib.2010.05.016 (2010).

4 Choi, R. et al. Taming the Boys for Global Good: Contraceptive Strategy to Stop Malaria Transmission. Molecules 25, doi:10.3390/molecules25122773 (2020).

5 Lawrence, C. J. et al. A standardized kinesin nomenclature. J Cell Biol 167, 19–22, doi:10.1083/jcb.200408113 (2004).

6 Depoix, D. et al. Vital role for Plasmodium berghei Kinesin8B in axoneme assembly during male gamete formation and mosquito transmission. Cell Microbiol 22, e13121, doi:10.1111/cmi.13121 (2020).

7 Zeeshan, M. et al. Kinesin-8B controls basal body function and flagellum formation and is key to malaria transmission. Life Sci Alliance 2, doi:10.26508/lsa.201900488 (2019).

8 Wickstead, B., Gull, K. & Richards, T. A. Patterns of kinesin evolution reveal a complex ancestral eukaryote with a multifunctional cytoskeleton. BMC Evol Biol 10, 110, doi:10.1186/1471-2148-10-110 (2010).

9 Walczak, C. E., Gayek, S. & Ohi, R. Microtubule-depolymerizing kinesins. Annu Rev Cell Dev Biol 29, 417–441, doi:10.1146/annurev-cellbio-101512-122345 (2013).

10 Zeeshan, M. et al. Plasmodium kinesin-8X associates with mitotic spindles and is essential for oocyst development during parasite proliferation and transmission. PLoS Pathog 15, e1008048, doi:10.1371/journal.ppat.1008048 (2019).

11 Arellano-Santoyo, H. et al. A Tubulin Binding Switch Underlies Kip3/Kinesin-8 Depolymerase Activity. Dev Cell 42, 37–51 e38, doi:10.1016/j.devcel.2017.06.011 (2017).

12 Mayr, M. I. et al. The human kinesin Kif18A is a motile microtubule depolymerase essential for chromosome congression. Curr Biol 17, 488–498, doi:10.1016/j.cub.2007.02.036 (2007).

13 Varga, V. et al. Yeast kinesin-8 depolymerizes microtubules in a length-dependent manner. Nat Cell Biol 8, 957–962, doi:10.1038/ncb1462 (2006).

14 Unsworth, A., Masuda, H., Dhut, S. & Toda, T. J. M. b. o. t. c. Fission yeast kinesin-8 Klp5 and Klp6 are interdependent for mitotic nuclear retention and required for proper microtubule dynamics. 19, 5104–5115 (2008).

15 Du, Y., English, C. A. & Ohi, R. The kinesin-8 Kif18A dampens microtubule plus-end dynamics. Curr Biol 20, 374–380, doi:10.1016/j.cub.2009.12.049 (2010).

16 Gardner, M. K., Odde, D. J. & Bloom, K. Kinesin-8 molecular motors: putting the brakes on chromosome oscillations. Trends Cell Biol 18, 307–310, doi:10.1016/j.tcb.2008.05.003 (2008).

17 Peters, C. et al. Insight into the molecular mechanism of the multitasking kinesin-8 motor. EMBO J 29, 3437–3447, doi:10.1038/emboj.2010.220 (2010).

18 Locke, J. et al. Structural basis of human kinesin-8 function and inhibition. Proc Natl Acad Sci U S A 114, E9539–E9548, doi:10.1073/pnas.1712169114 (2017).

19 Wang, D. et al. Motility and microtubule depolymerization mechanisms of the Kinesin-8 motor, KIF19A. Elife 5, doi:10.7554/eLife.18101 (2016).

20 Wang, W. et al. New Insights into the Coupling between Microtubule Depolymerization and ATP Hydrolysis by Kinesin-13 Protein Kif2C. J Biol Chem 290, 18721–18731, doi:10.1074/jbc.M115.646919 (2015).

21 Rice, S. et al. A structural change in the kinesin motor protein that drives motility. Nature 402, 778–784, doi:10.1038/45483 (1999).

22 Atherton, J. et al. The divergent mitotic kinesin MKLP2 exhibits atypical structure and mechanochemistry. Elife 6, doi:10.7554/eLife.27793 (2017).

23 Atherton, J. et al. Conserved mechanisms of microtubule-stimulated ADP release, ATP binding, and force generation in transport kinesins. Elife 3, e03680, doi:10.7554/eLife.03680 (2014).

24 Shang, Z. et al. High-resolution structures of kinesin on microtubules provide a basis for nucleotide-gated force-generation. Elife 3, e04686, doi:10.7554/eLife.04686 (2014).

25 Cook, A. D. et al. Cryo-EM structure of a microtubule-bound parasite kinesin motor and implications for its mechanism and inhibition. J Biol Chem 297, 101063, doi:10.1016/j.jbc.2021.101063 (2021).

26 Benoit, M. et al. Structural basis of mechano-chemical coupling by the mitotic kinesin KIF14. Nat Commun 12, 3637, doi:10.1038/s41467-021-23581-3 (2021).

27 Woehlke, G. et al. Microtubule interaction site of the kinesin motor. Cell 90, 207–216 (1997).

28 Hirokawa, N., Nitta, R. & Okada, Y. The mechanisms of kinesin motor motility: lessons from the monomeric motor KIF1A. Nat Rev Mol Cell Biol 10, 877–884, doi:10.1038/nrm2807 (2009).

29 Erent, M., Drummond, D. R. & Cross, R. A. S. pombe kinesins-8 promote both nucleation and catastrophe of microtubules. PLoS One 7, e30738, doi:10.1371/journal.pone.0030738 (2012).

30 Cao, L. et al. The structure of apo-kinesin bound to tubulin links the nucleotide cycle to movement. Nat Commun 5, 5364, doi:10.1038/ncomms6364 (2014).

31 Vale, R. D. & Milligan, R. A. The way things move: looking under the hood of molecular motor proteins. Science 288, 88–95, doi:10.1126/science.288.5463.88 (2000).

32 Arellano-Santoyo, H. et al. Multimodal tubulin binding by the yeast kinesin-8, Kip3, underlies its motility and depolymerization. bioRxiv, 2021.2010.2012.464151, doi:10.1101/2021.10.12.464151 (2021).

33 Kim, H., Fonseca, C. & Stumpff, J. A unique kinesin-8 surface loop provides specificity for chromosome alignment. Mol Biol Cell 25, 3319–3329, doi:10.1091/mbc.E14-06-1132 (2014).

34 Parke, C. L., Wojcik, E. J., Kim, S. & Worthylake, D. K. ATP hydrolysis in Eg5 kinesin involves a catalytic two-water mechanism. J Biol Chem 285, 5859–5867, doi:10.1074/jbc.M109.071233 (2010).

35 Benoit, M. P. M. H., Asenjo, A. B. & Sosa, H. Cryo-EM reveals the structural basis of microtubule depolymerization by kinesin-13s. Nat Commun 9, 1662, doi:10.1038/s41467-018-04044-8 (2018).

36 Ems-McClung, S. C. & Walczak, C. E. Kinesin-13s in mitosis: Key players in the spatial and temporal organization of spindle microtubules. Semin Cell Dev Biol 21, 276–282, doi:10.1016/j.semcdb.2010.01.016 (2010).

37 Friel, C. T. & Welburn, J. P. Parts list for a microtubule depolymerising kinesin. Biochem Soc Trans 46, 1665–1672, doi:10.1042/BST20180350 (2018).

38 Shipley, K. et al. Structure of a kinesin microtubule depolymerization machine. EMBO J 23, 1422–1432, doi:10.1038/sj.emboj.7600165 (2004).

39 Ogawa, T., Saijo, S., Shimizu, N., Jiang, X. & Hirokawa, N. Mechanism of Catalytic Microtubule Depolymerization via KIF2-Tubulin Transitional Conformation. Cell Rep 20, 2626–2638, doi:10.1016/j.celrep.2017.08.067 (2017).

40 Hirst, W. G. et al. Purification of functional Plasmodium falciparum tubulin allows for the identification of parasite-specific microtubule inhibitors. Curr Biol, doi:10.1016/j.cub.2021.12.049 (2022).

41 Invergo, B. M. et al. Sub-minute Phosphoregulation of Cell Cycle Systems during Plasmodium Gamete Formation. Cell Rep 21, 2017–2029, doi:10.1016/j.celrep.2017.10.071 (2017).

42 Bushell, E. et al. Functional Profiling of a Plasmodium Genome Reveals an Abundance of Essential Genes. Cell 170, 260-272.e268, doi:10.1016/j.cell.2017.06.030 (2017).

43 Lasonder, E. et al. Integrated transcriptomic and proteomic analyses of P. falciparum gametocytes: molecular insight into sex-specific processes and translational repression. Nucleic Acids Res 44, 6087–6101, doi:10.1093/nar/gkw536 (2016).

44 Avidor-Reiss, T. & Leroux, M. R. Shared and Distinct Mechanisms of Compartmentalized and Cytosolic Ciliogenesis. Curr Biol 25, R1143–1150, doi:10.1016/j.cub.2015.11.001 (2015).

45 Sinden, R. E., Canning, E. U. & Spain, B. Gametogenesis and fertilization in Plasmodium yoelii nigeriensis: a transmission electron microscope study. Proc R Soc Lond B Biol Sci 193, 55–76, doi:10.1098/rspb.1976.0031 (1976).

46 Persico, V., Callaini, G. & Riparbelli, M. G. The Microtubule-Depolymerizing Kinesin-13 Klp10A Is Enriched in the Transition Zone of the Ciliary Structures of. Front Cell Dev Biol 7, 173, doi:10.3389/fcell.2019.00173 (2019).

47 Blaineau, C. et al. A novel microtubule-depolymerizing kinesin involved in length control of a eukaryotic flagellum. Curr Biol 17, 778–782, doi:10.1016/j.cub.2007.03.048 (2007).

48 Chan, K. Y., Matthews, K. R. & Ersfeld, K. Functional characterisation and drug target validation of a mitotic kinesin-13 in Trypanosoma brucei. PLoS Pathog 6, e1001050, doi:10.1371/journal.ppat.1001050 (2010).

49 Dawson, S. C. et al. Kinesin-13 regulates flagellar, interphase, and mitotic microtubule dynamics in Giardia intestinalis. Eukaryot Cell 6, 2354–2364, doi:10.1128/EC.00128-07 (2007).

50 McInally, S. G., Kondev, J. & Dawson, S. C. Length-dependent disassembly maintains four different flagellar lengths in. Elife 8, doi:10.7554/eLife.48694 (2019).

51 Niwa, S. et al. KIF19A is a microtubule-depolymerizing kinesin for ciliary length control. Dev Cell 23, 1167–1175, doi:10.1016/j.devcel.2012.10.016 (2012).

52 Piao, T. et al. A microtubule depolymerizing kinesin functions during both flagellar disassembly and flagellar assembly in Chlamydomonas. Proc Natl Acad Sci U S A 106, 4713–4718, doi:10.1073/pnas.0808671106 (2009).

53 Zeeshan, M. et al. Location and function of Plasmodium kinesins: key roles in parasite proliferation, polarity, and transmission. bioRxiv, 2021.2005.2026.445751, doi:10.1101/2021.05.26.445751 (2021).

54 De Marco, V., Stier, G., Blandin, S. & de Marco, A. The solubility and stability of recombinant proteins are increased by their fusion to NusA. Biochem Biophys Res Commun 322, 766–771, doi:10.1016/j.bbrc.2004.07.189 (2004).

55 Kreuzer, K. N. & Jongeneel, C. V. Escherichia coli phage T4 topoisomerase. Methods Enzymol 100, 144–160, doi:10.1016/0076-6879(83)00051-8 (1983).

56 Toropova, K., Mladenov, M. & Roberts, A. J. Intraflagellar transport dynein is autoinhibited by trapping of its mechanical and track-binding elements. Nat Struct Mol Biol 24, 461–468, doi:10.1038/nsmb.3391 (2017).

57 Schindelin, J. et al. Fiji: an open-source platform for biological-image analysis. Nat Methods 9, 676–682, doi:10.1038/nmeth.2019 (2012).

58 Mastronarde, D. N. Automated electron microscope tomography using robust prediction of specimen movements. J Struct Biol 152, 36–51, doi:10.1016/j.jsb.2005.07.007 (2005).

59 Rohou, A. & Grigorieff, N. CTFFIND4: Fast and accurate defocus estimation from electron micrographs. J Struct Biol 192, 216–221, doi:10.1016/j.jsb.2015.08.008 (2015).

60 Tang, G. et al. EMAN2: an extensible image processing suite for electron microscopy. J Struct Biol 157, 38–46, doi:10.1016/j.jsb.2006.05.009 (2007).

61 Cook, A. D., Manka, S. W., Wang, S., Moores, C. A. & Atherton, J. A microtubule RELION-based pipeline for cryo-EM image processing. J Struct Biol 209, 107402, doi:10.1016/j.jsb.2019.10.004 (2020).

62 Zivanov, J. et al. New tools for automated high-resolution cryo-EM structure determination in RELION-3. Elife 7, doi:10.7554/eLife.42166 (2018).

63 Zivanov, J., Nakane, T. & Scheres, S. H. W. Estimation of high-order aberrations and anisotropic magnification from cryo-EM data sets in RELION-3.1. IUCrJ 7, 253–267, doi:10.1107/S2052252520000081 (2020).

64 Punjani, A., Rubinstein, J. L., Fleet, D. J. & Brubaker, M. A. cryoSPARC: algorithms for rapid unsupervised cryo-EM structure determination. Nat Methods 14, 290–296, doi:10.1038/nmeth.4169 (2017).

65 Sali, A. & Blundell, T. L. Comparative protein modelling by satisfaction of spatial restraints. J Mol Biol 234, 779–815, doi:10.1006/jmbi.1993.1626 (1993).

66 Arora, K. et al. KIF14 binds tightly to microtubules and adopts a rigor-like conformation. J Mol Biol 426, 2997–3015, doi:10.1016/j.jmb.2014.05.030 (2014).

67 Dong, G. Q., Fan, H., Schneidman-Duhovny, D., Webb, B. & Sali, A. Optimized atomic statistical potentials: assessment of protein interfaces and loops. Bioinformatics 29, 3158–3166, doi:10.1093/bioinformatics/btt560 (2013).

68 Benkert, P., Künzli, M. & Schwede, T. QMEAN server for protein model quality estimation. Nucleic Acids Res 37, W510–514, doi:10.1093/nar/gkp322 (2009).

69 Goddard, T. D., Huang, C. C. & Ferrin, T. E. Visualizing density maps with UCSF Chimera. J Struct Biol 157, 281–287, doi:10.1016/j.jsb.2006.06.010 (2007).

70 Wang, R. Y. et al. Automated structure refinement of macromolecular assemblies from cryo-EM maps using Rosetta. Elife 5, doi:10.7554/eLife.17219 (2016).

71 Wang, C., Bradley, P. & Baker, D. Protein-protein docking with backbone flexibility. J Mol Biol 373, 503–519, doi:10.1016/j.jmb.2007.07.050 (2007).

72 Mandell, D. J., Coutsias, E. A. & Kortemme, T. Sub-angstrom accuracy in protein loop reconstruction by robotics-inspired conformational sampling. Nat Methods 6, 551–552, doi:10.1038/nmeth0809-551 (2009).

73 Madeira, F. et al. The EMBL-EBI search and sequence analysis tools APIs in 2019. Nucleic Acids Res 47, W636–W641, doi:10.1093/nar/gkz268 (2019).

74 Larkin, M. A. et al. Clustal W and Clustal X version 2.0. Bioinformatics 23, 2947–2948, doi:10.1093/bioinformatics/btm404 (2007).

75 Cross, R. A. Review: Mechanochemistry of the kinesin-1 ATPase. Biopolymers 105, 476–482, doi:10.1002/bip.22862 (2016).

