## Supplementary Data for "Mechanochemical tuning of a kinesin motor essential for malaria parasite transmission"

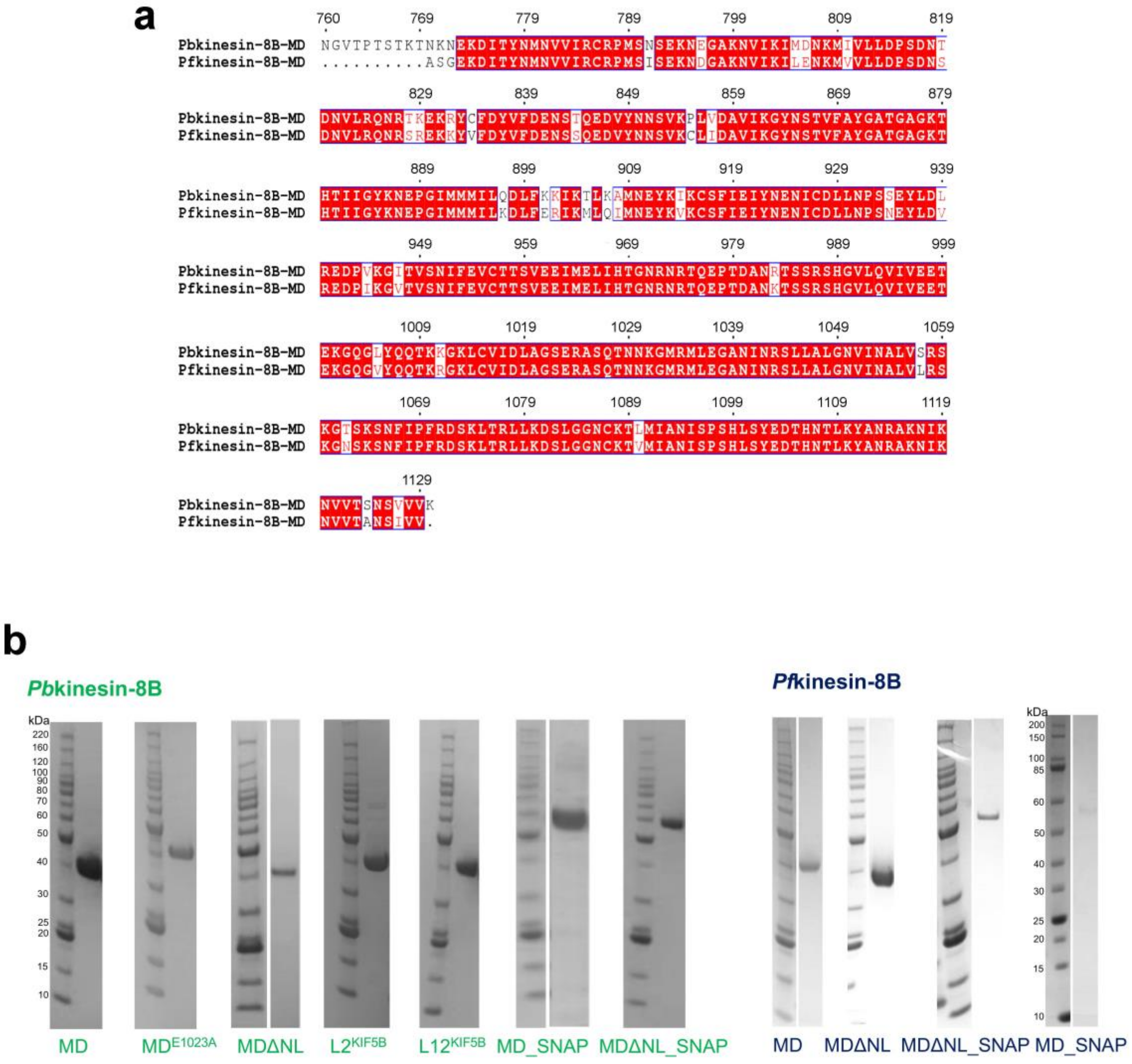


**Supplementary Figure 1. Sequence alignment of *Pb*kinesin-8B-MD with *Pf*kinesin-8B-MD and purified kinesin-8B proteins.**

a) EMBOSS needle alignment shows 88% identity (shaded red) and 94% similarity (red text) between *Pb*kinesin-8B-MD with *Pf*kinesin-8B-MD protein sequences.

b) SDS-PAGE Coomassie stained gels showing pure proteins used in our experiments. BenchMark™ Protein Ladder (Invitrogen) was used for all gels apart from the one for *Pf*kinesin-8B-MD-SNAP, for which unstained Protein Standard (Broad Range, NEB) was used. Empty space between sample lane and marker indicates these 2 lanes are from the same gel but not next to each other.

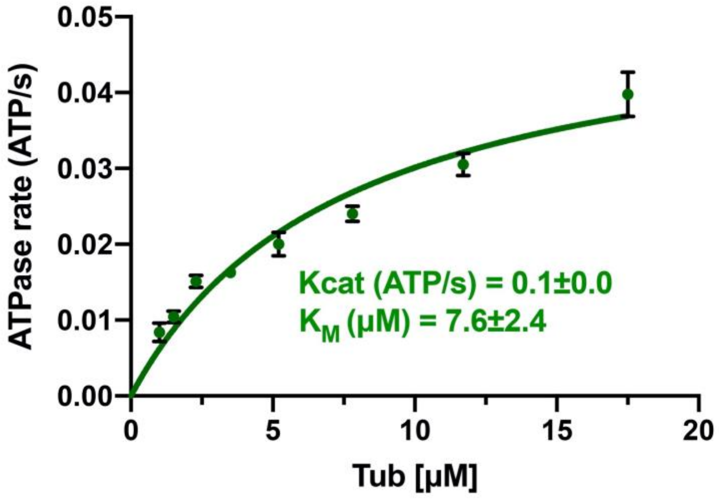


**Supplementary Figure 2. *Pb*kinesin-8B-MD exhibits tubulin stimulated ATPase activity.** The ATPase assay data (n=3 for each point, mean ± SD) were fitted using Michaelis-Menten equation, from which the Kcat and $K_M$ were calculated in Prism9.

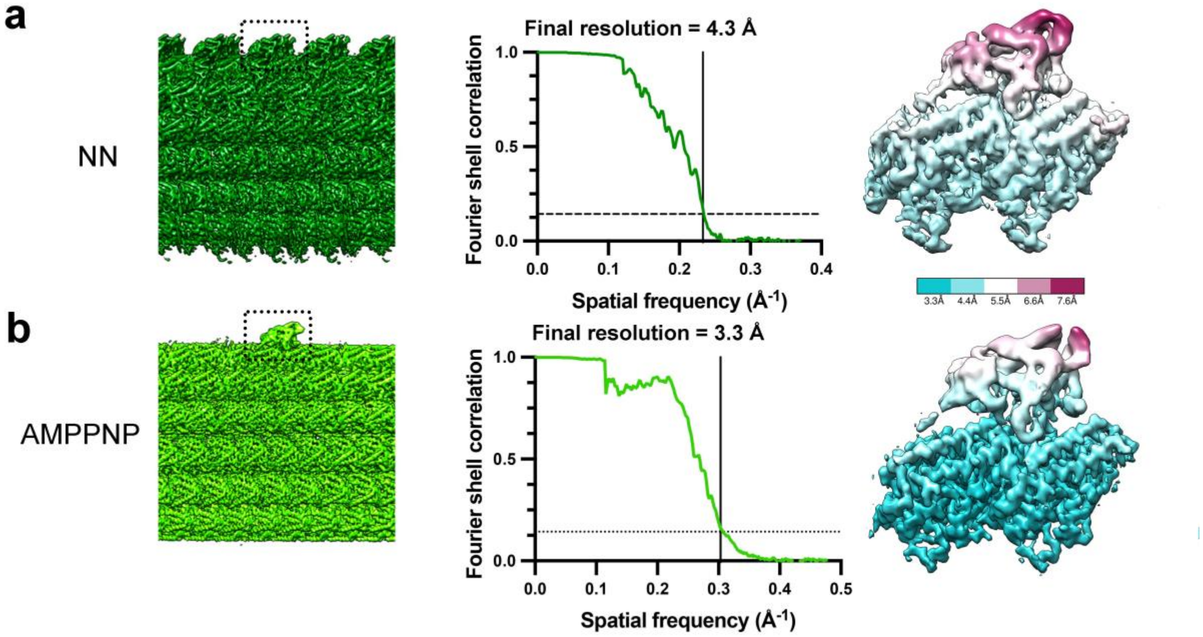


**Supplementary Figure 3. Cryo-EM reconstruction resolution estimation.**

a) Left, final reconstruction for *Pb*kinesin-8B-MD bound MT (dotted rectangle) in the absence of nucleotide following 3D classification to optimise motor domain density; in this case, the top protofilament is enriched for the motor domain; data processing details are described in the Methods; middle, global resolution FSC curve; right, local resolution for best kinesin motor-tubulin dimer.

b) Left, final reconstruction for *Pb*kinesin-8B-MD bound MT (dotted rectangle) in the presence of AMPPNP following 3D classification to optimise motor domain density; in this case, the central motor-tubulin dimer asymmetric unit is enriched; data processing details are described in the Methods; middle, global resolution FSC curve; right, local resolution for best kinesin motor-tubulin dimer.

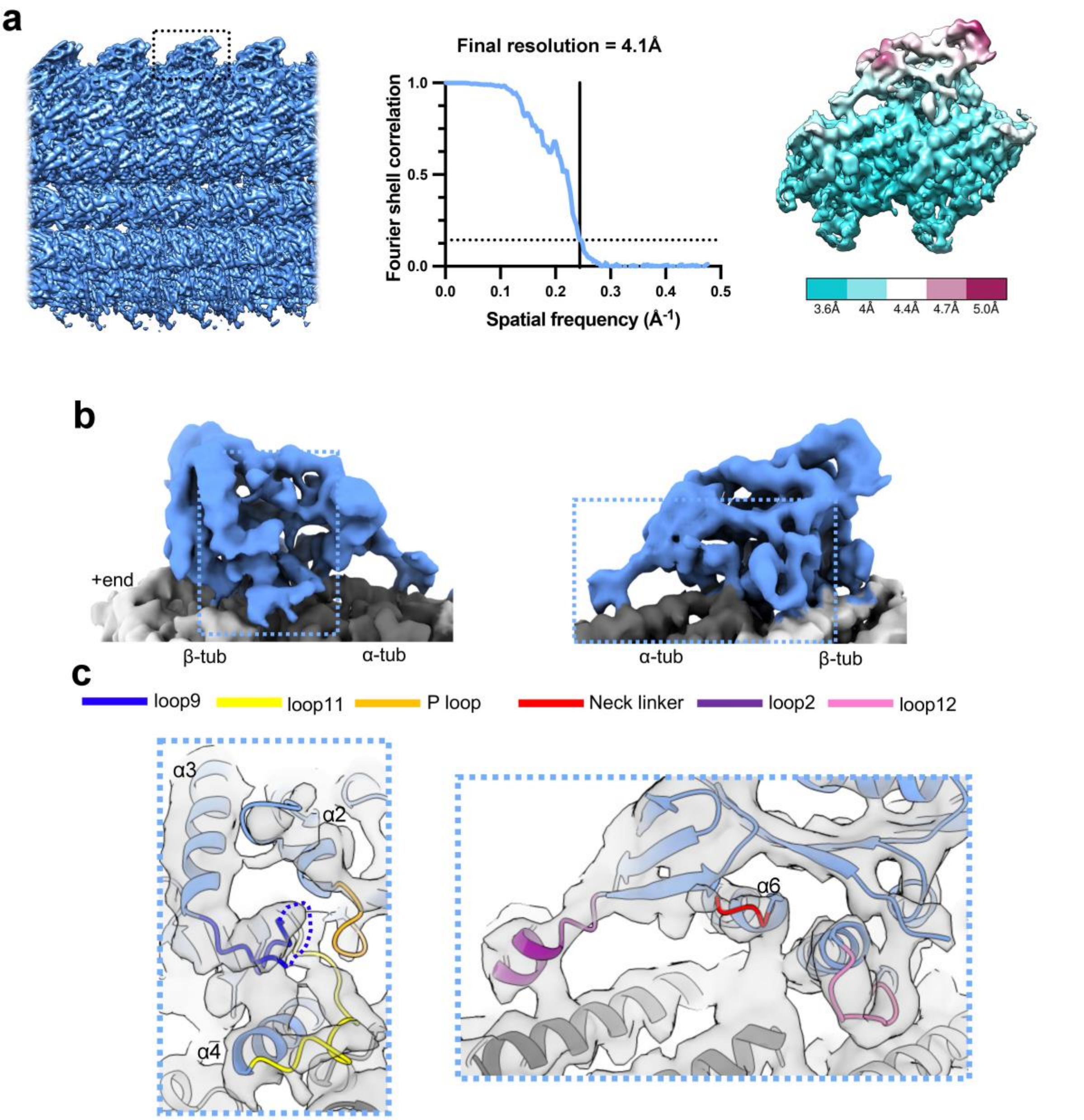


**Supplementary Figure 4. Cryo-EM reconstruction of MT-bound *Pf*kinesin-8B-MD NN and resolution estimation.**

a) Left, final reconstruction for *Pf*kinesin-8B-MD bound MT (dotted rectangle) in the absence of nucleotide following 3D classification; middle, global resolution FSC curve; right, local resolution density for best kinesin motor-tubulin dimer.

b) Asymmetric unit of MT-bound NN *Pf*kinesin-8B-MD depicted as solid surface representation and viewed towards the NBS (left) and the neck linker region (right). *Pf*kinesin-8B-MD-NN density is coloured in blue, α- and β-tubulin are coloured in dark and light grey, respectively; regions around NBS and neck linker depicted in (c) are boxed.

c) Left, zoom-in view of the NBS of NN *Pf*kinesin-8B-MD with docked model; right: zoom-in view of the neck linker region of NN *Pf*kinesin-8B-MD with docked model. *Pf*kinesin-8B-MD-NN model is coloured in blue and α- and β-tubulin are coloured in dark and light grey, respectively.

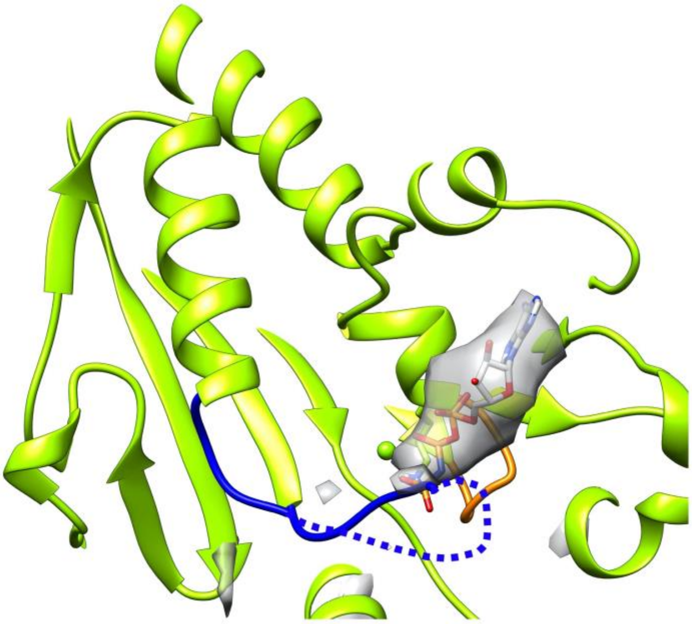

**Supplementary Figure 5. AMPPNP binds to MT-bound *Pb*kinesin-8B-MD.**
Difference density calculated between *Pb*kinesin-8B-MD-AMPPNP reconstruction and simulated 7 Å resolution density of the protein-only model. The difference map was calculated using Chimera “vop subtract” command. Density shown in transparent grey superimposed on the *Pb*kinesin-8B-MD-AMPPNP model, and corresponding with AMPPNP bound at the NBS. AMPPNP *Pb*kinesin-8B-MD model is coloured in light green, the P-loop is orange and flexible-appearing loop 9 is the dashed blue line.

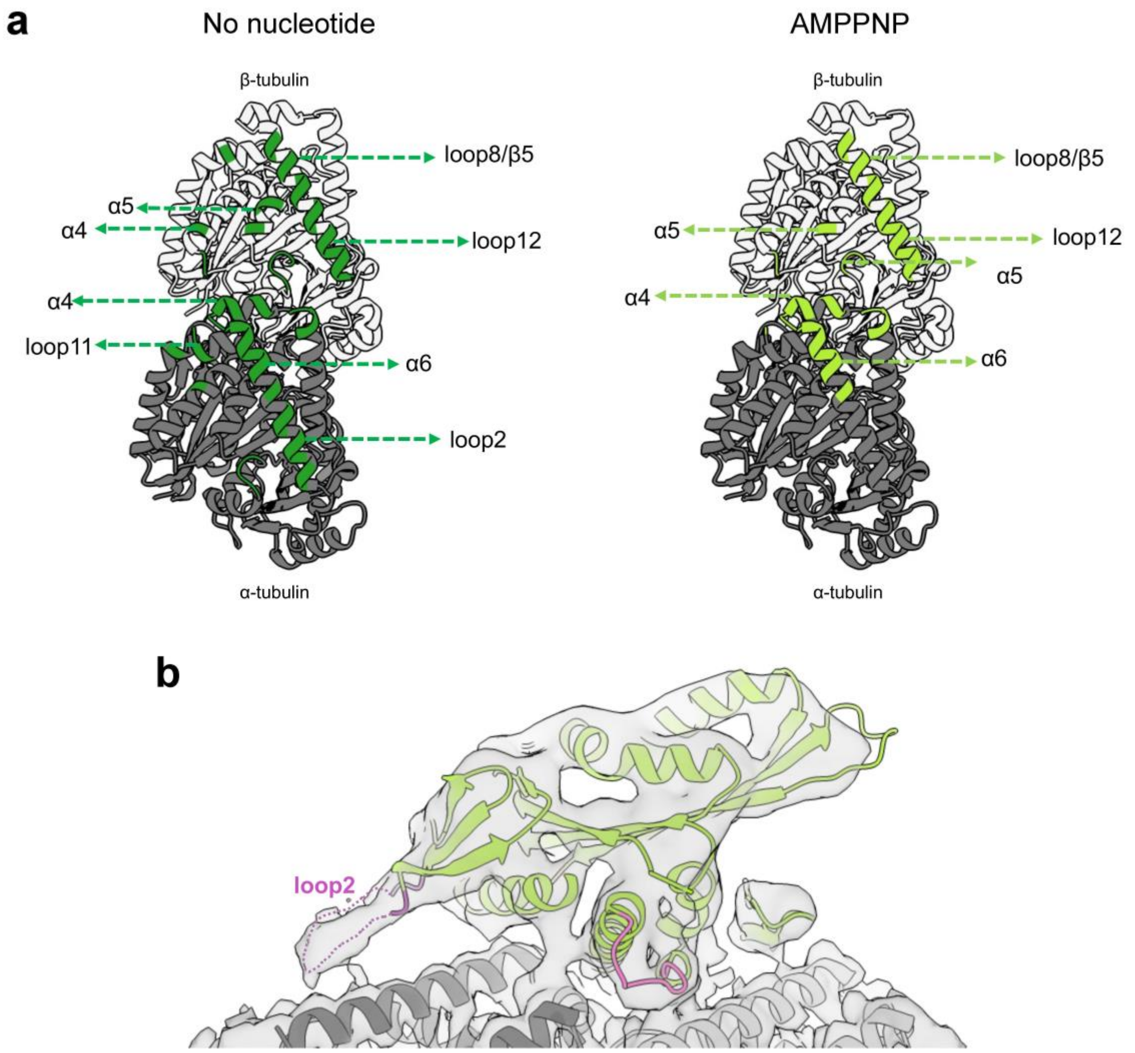


**Supplementary Figure 6. Comparison of nucleotide-dependent *Pb*kinesin-8B-MD footprints on the MT surface.**

a) Contacts formed between *Pb*kinesin-8B-MD and the MT in NN state and AMPPNP state are very similar. Left: MT footprint of *Pb*kinesin-8B-MD in NN state in dark green; dashed lines indicate contacting secondary structure elements in *Pb*kinesin-8B-MD. Tubulin residues <5Å distance from the bound motor is colored in dark green. Right: MT footprint of *Pb*kinesin-8B-MD in AMPPNP state in yellow green. α-tubulin are otherwise depicted in dark grey ribbon and β-tubulin in light grey ribbon.

b) Lower threshold view of the of MT-bound *Pb*kinesin-8B-MD in the AMPPNP state showing the contact formed between $\alpha$-tubulin and loop2 (purple).

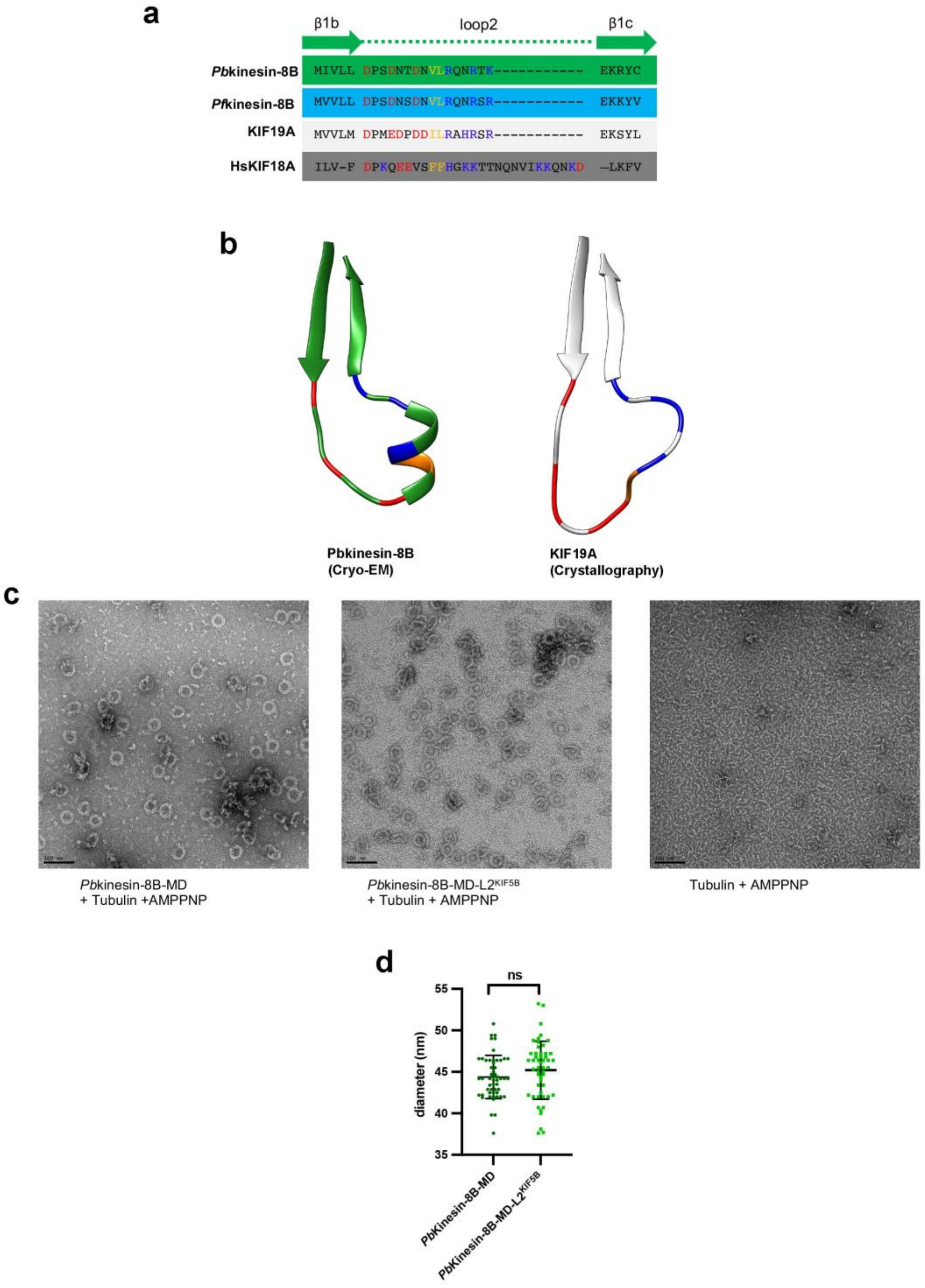


**Supplementary Figure 7. Comparison of loop 2 of mammalian and malaria kinesin-8s, and characterisation of *Pb*kinesin-8B-MD loop 2 mutant-induced tubulin ring-like structures.**

a) Sequence alignment of loop 2 from *Pb*kinesin-8B, *Pf*kinesin-8B, human KIF18A and mouse KIF19A. Positively charged residues are coloured blue, negatively charged residues are coloured red and hydrophobic residues are coloured orange.

b) Loop 2 of *Pb*kinesin-8B (MT-bound NN model) and KIF19A (PDB 5GSZ) are the same length but adopt different conformations.

c) Negative stain EM images showing the formation of tubulin rings by incubation of WT *Pb*kinesin-8B-MD or *Pb*kinesin-8B-MD-L2$^{KIF5B}$ with tubulin and AMPPNP; these structures do not form in the absence of *Pb*kinesin-8B. Scale bar: 100nm.

d) The diameter of tubulin rings formed by *Pb*kinesin-8B-MD-L2$^{KIF5B}$ is indistinguishable from those formed by WT *Pb*kinesin-8B-MD. Mean diameter of rings formed by WT *Pb*kinesin-8B-MD = 44.4 ± 2.6nm(mean ± SD, n = 53); mean diameter of rings formed by *Pb*kinesin-8B-MD-L2$^{KIF5B}$ = 45.2 ± 3.5nm (mean ± SD, n = 51). ns, not significant. p =0.1745 by t-test.

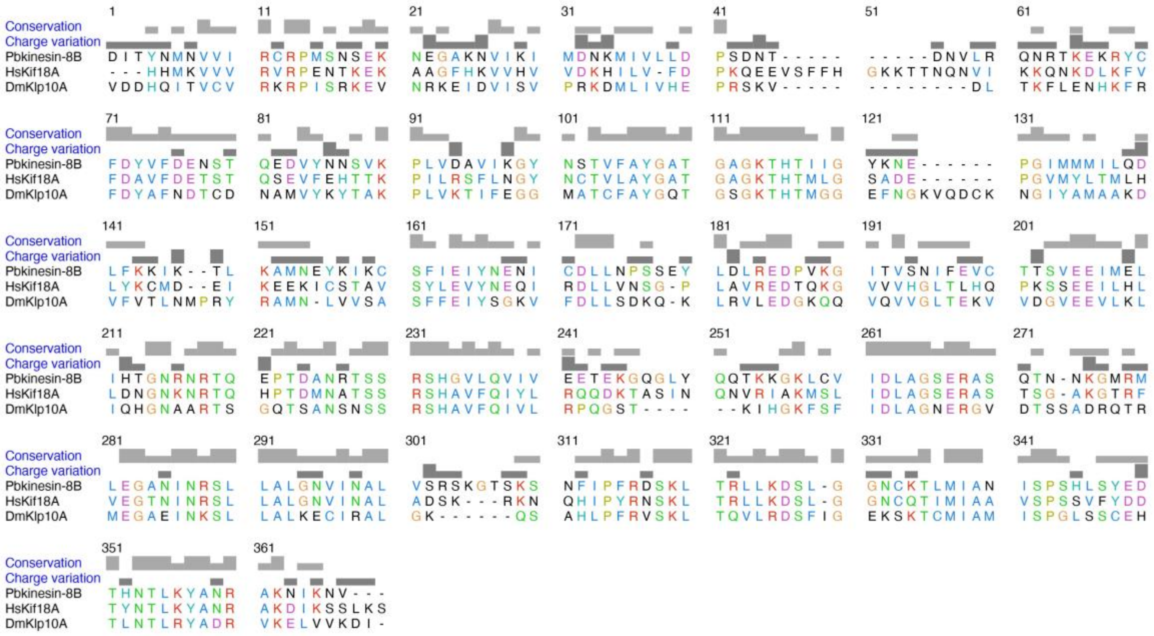

**Supplementary Figure 8. Sequence alignments of *Pb*kinesin-8B, HsKif18A and DmKlp10A motor domains.**

Alignments were done with Clustal Omega. Residue colouring uses the Clustal X scheme.
